# Osmotic-stress-inducible nuclear condensates restrict gene inducibility

**DOI:** 10.64898/2026.06.01.729169

**Authors:** Hikaru Sato, Satoru Fujimoto, Marie Sakuma, Miki Fujita, Daniel Slane, Emi Mishiro-Sato, Emi Yumoto, Masashi Asahina, Akinori Kanai, Yutaka Suzuki, Fuminori Takahashi, Kazuko Yamagushi-Shinozaki, Kazuo Shinozaki, Sachihiro Matsunaga

## Abstract

Plants, as sessile organisms, have developed various mechanisms to respond to environmental stress conditions. The plant hormone abscisic acid (ABA) is necessary for the plant to adapt to osmotic stress conditions. However, the molecular mechanisms preceding ABA accumulation remain largely unknown. To isolate transcriptional complexes on the promoter region of *NINE-CIS-EPOXYCAROTENOID DIOXYGENASE 3* (*NCED3*) encoding a rate-limiting enzyme in the ABA biosynthetic pathway *in planta*, we developed the insertional chromatin immunoprecipitation (iChIP) screen method. The identified ALBA proteins formed condensates through liquid-liquid phase separation (LLPS) in response to osmotic stress conditions. ALBA4 directly binds to stress-inducible genes, including *NCED3*, and suppresses their stress inducibility. Our results demonstrate how plants respond to osmotic stress at early timepoints before ABA biosynthesis through condensate formation as osmo-sensors.

## Main Text

Plants have developed various strategies to respond to environmental changes as sessile organisms. Water-deficit stress conditions, including drought and high salt stresses, frequently occur in natural environments, especially during the currently ongoing global warming (*1, 2*). The water-deficit conditions cause osmotic stresses for plants, and they severely suppress plant growth and reproduction resulting in loss of agricultural products. Physiologically, osmotic stress inhibits photosynthesis and transpiration, which induce higher leaf temperature (*3, 4*). At a cellular level, osmotic conditions decrease turgor pressure with decreased cell sizes and destabilize membrane structures and proteins (*5, 6*). Abscisic acid (ABA) is a plant hormone that is accumulated under stress conditions, and has critical roles for plants to adapt to water-deficit conditions (*7*) through stomatal closure, accumulation of osmo-protectants, and gene expression of stress-inducible genes (*8*). Previous studies have revealed the detailed molecular mechanisms by which ABA is transported (*9*) and perceived (*10*) to activate the downstream signaling cascades (*11*). However, molecular mechanisms at earlier timepoints before ABA accumulation remain largely unknown.

In the ABA biosynthetic pathway, NINE-CIS-EPOXYCAROTENOID DIOXYGENASE family proteins are known as rate-limiting enzymes on the basis of their stress inducibility, and transcriptional induction of *NCED3* plays a dominant role in stress-induced ABA accumulation among the *NCED* family genes in *Arabidopsis thaliana* (Arabidopsis). The single *nced3* knockout mutant showed severe drought stress sensitivity (*12*). Several transcription factors and epigenetic factors have been reported to regulate gene expression of *NCED3* (*13–15*). However, the detailed molecular mechanisms of how the *NCED3* gene expression is regulated remain elusive.

Recently, the formation of biological molecular condensates through liquid-liquid phase separation (LLPS) has been reported as a sensing mechanism of environmental changes. Formation of biomolecular condensates is directly triggered by changes in environmental physical and chemical conditions, such as osmotic, heat, and cold stresses, and it involves various molecular mechanisms, including transcriptional and post-transcriptional regulation (*16, 17*). In Arabidopsis, a transcription factor SEUSS (SEU) (*18*) and a mRNA decapping enzyme DECAPPING 5 (DCP5) (*19*) have been reported as potential osmo-sensing proteins that form condensates through LLPS by sensing cellular molecular crowding induced by cell shrinkage under osmotic conditions (*20*). However, detailed mechanisms of whether and how the formation of condensates affects ABA biosynthesis have not yet been well explored.

Here we applied insertional chromatin immunoprecipitation (iChIP) (*21*) to isolate a transcriptional complex on the *NCED3* promoter *in planta*. As a result, ACETYLATION LOWERS BINDING AFFINITY (ALBA) proteins were identified. The ALBA proteins directly bound to and suppressed the transcriptional inducibility of osmotic stress-inducible genes, including *NCED3*, and formed osmotic-stress inducible condensates through LLPS, which serves as potential osmo-sensors by molecular crowding.

### Identification of ALBA proteins by iChIP

To identify a transcriptional complex on the *NCED3* promoter *in planta*, we applied an insertional chromatin immunoprecipitation (iChIP) screen method for plants which had been developed in mammalian cells (*21*). In the iChIP screen, artificial tandem repeated elements are fused to a specific promoter region and the binding protein is expressed (Fig. 1, A and B). The transcriptional complex on the promoter can be immunoprecipitated through a protein tag fused to the binding protein (Fig. 1C). For the optimized iChIP for plants in this study, we used the GAL4-binding domain (GAL4-BD) and the upstream activating sequences (UAS) which harbor the binding elements of GAL4-BD. The repeated UAS motifs were fused to the 2.3-kb promoter of *NCED3*, which has been reported to be required for the gene activation (*22*). Additionally, to quantitatively evaluate the result, negative control plants without the repeated UAS were generated (Fig. 1B). After confirming the reporter *GUS* gene was induced by dehydration stress (Fig. 1D) and that the GAL4-BD-GFP-nls fusion protein bound to the UAS (Fig. 1E), two replicates of iChIP were performed during dehydration stress (Fig. 1F and table S1). Among the reproducibly enriched candidates, we focused on ALBA4 (At1g76010), ALBA5 (AT1G20220) and ALBA6 (At3g07030) (ALBA4/5/6) (Fig. 1G) as all family proteins in the same subclade were identified and because ALBA orthologous proteins are reported to function in nuclei (*23, 24*). According to gene expression analyses of *ALBA4/5/6* and GUS staining experiments using their respective promoters, we found that the gene expression of *ALBA4/5/6* was stable during dehydration and salt stresses (fig. S1, A and B). Additionally, their tissue-specific expression patterns were not largely changed under the stress conditions (fig. S1, C to E). Additionally, the *ALBA6* promoter was activated in distinct tissues such as shoot meristematic regions (fig. S1E), suggesting that *ALBA4* and *ALBA5* should have major functions in most tissues during osmotic stress.

**Fig. 1.**
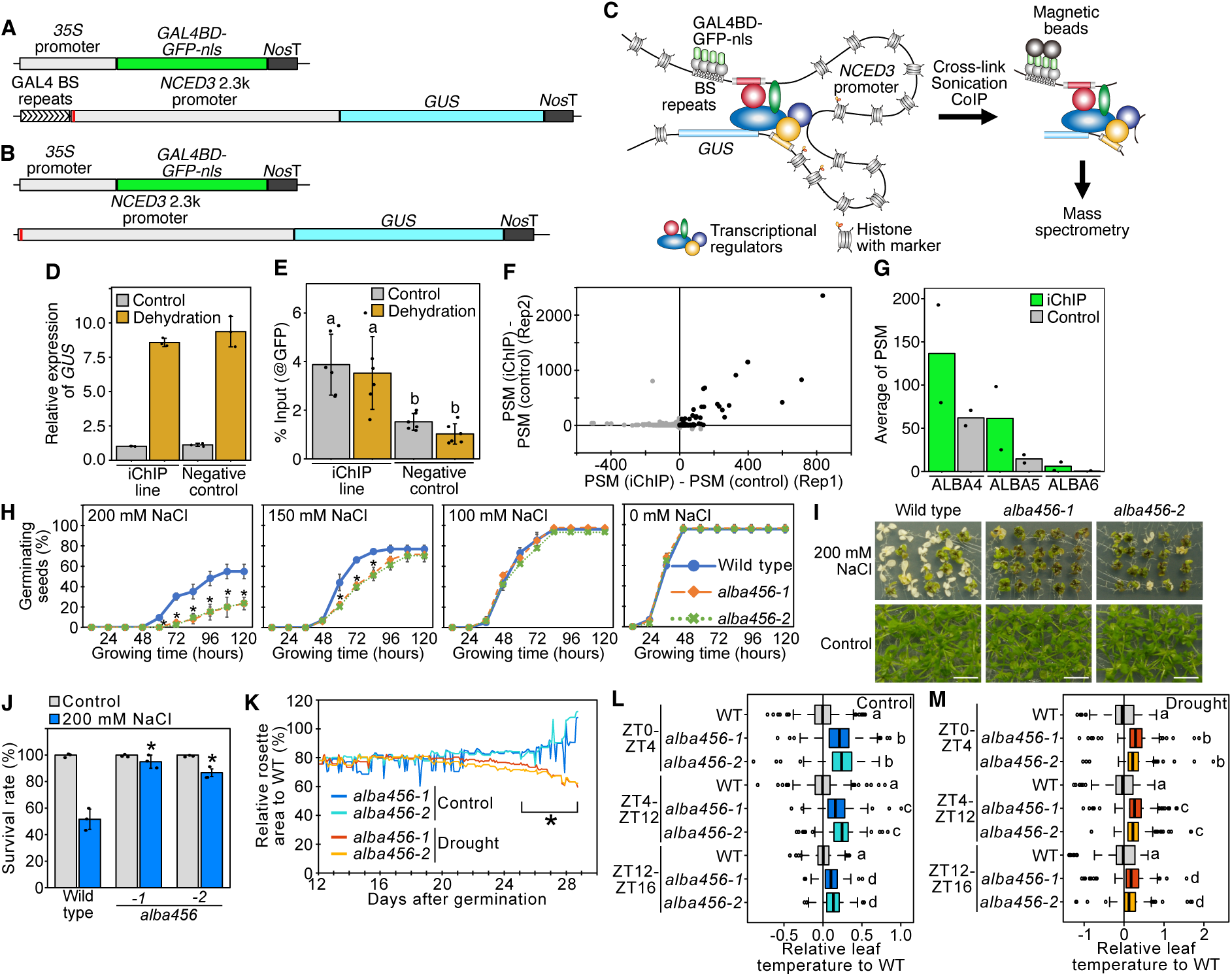
Identification of ALBAs as novel candidate regulators of *NCED3* during osmotic stress. (**A** and **B**) Schematic diagram of iChIP constructs (A) and negative control (B). (**C**) Schematic model of iChIP isolating a transcriptional complex on a promoter through the repeated motifs. (**D**) Expression levels of *GUS* in the iChIP and negative control lines. Error bars indicate SD from triplicate technical repeats. (**E**) ChIP assays of GAL4BD-GFP-nls proteins binding to the *NCED3* promoter region. Error bars indicate SD from total six technical replicates from two independent biological replicates. Letters above bars indicate significant differences (*P* < 0.05, Tukey’s multiple range test). (**F**) Scattered plot of identified proteins through two biological replicates of iChIP. (**G**) Average PSM of ALBA proteins in iChIP. (**H**) Germination ratios of *alba456* triple mutants under slat stress conditions. Error bars indicate SD from three biological replicates. Asterisks indicates significant differences in both mutant lines from wild type (*P* < 0.05, Tukey’s multiple range test). (**I** and **J**) Salt stress tolerance of *alba456* triple mutants. Images with and without salt stress (I) and the survival ratios (J) are shown. Error bars indicate SD from three biological replicates. Asterisks indicates significant differences from wild type (*P* < 0.05, Tukey’s multiple range test). (**K**) Drought stress sensitivity of *alba456* triple mutants. Relative rosette areas of the *alba456* triple mutants to wild type under control and drought stress are shown. The lines indicate average of 6 plants under each condition. Asterisks indicates significant differences between control and drought conditions in each line (*P* < 0.05, Student’s t test). (**L** and **M**) Leaf temperature in *alba456* triple mutants under control (L) and drought stress conditions (M). Relative leaf temperature to wild type during each circadian time period are shown. Letters above bars indicate significant differences (*P* < 0.05, Tukey’s multiple range test).

### ALBAs suppress osmotic stress responses

To assess how ALBA4/5/6 function under osmotic stress conditions, we generated two lines of triple mutants of *ALBA4*, *ALBA5*, and *ALBA6* (*alba456-1* and *alba456-2*) using the CRISPR Cas9 system (fig. S2, A to C). In germination assays, the *alba456* triple mutants exhibited significantly delayed germination (Fig. 1H). Additionally, the triple mutants showed higher salt stress tolerance with significantly higher survival ratios (Fig. 1, I and J). These triple mutant phenotypes were completely recovered by complementation with ALBA4-GFP and ALBA5-GFP driven by their native promoters (fig. S2, D to I). Next, to evaluate the phenotypes of the triple mutant under drought stress conditions, a RIKEN Integrated Plant Phenotyping System (RIPPS) analysis was conducted (*25*). Under well-watered and drought stress conditions (fig. 2J and table S2), rosette leaf areas and leaf temperature of the wild type and *alba456* mutants were measured continuously and automatically (fig. S2, K to P, and tables S3 and S4). The results showed that the *alba456* mutans had drought-sensitive phenotypes with more severe growth retardation and higher leaf temperature in comparison to wild type (Fig. 1, K to M). Especially, the increase in leaf temperature in *alba456* mutants was most pronounced during the early daytime period under both control and drought stress conditions (ZT0-4) (Fig. 1, L and M), suggesting circadian clock-dependent functions of ALBAs. Taken together, ALBA4/5/6 function to suppress osmotic stress responses in plants and balance plant growth and stress tolerance.

**Fig. 2.**
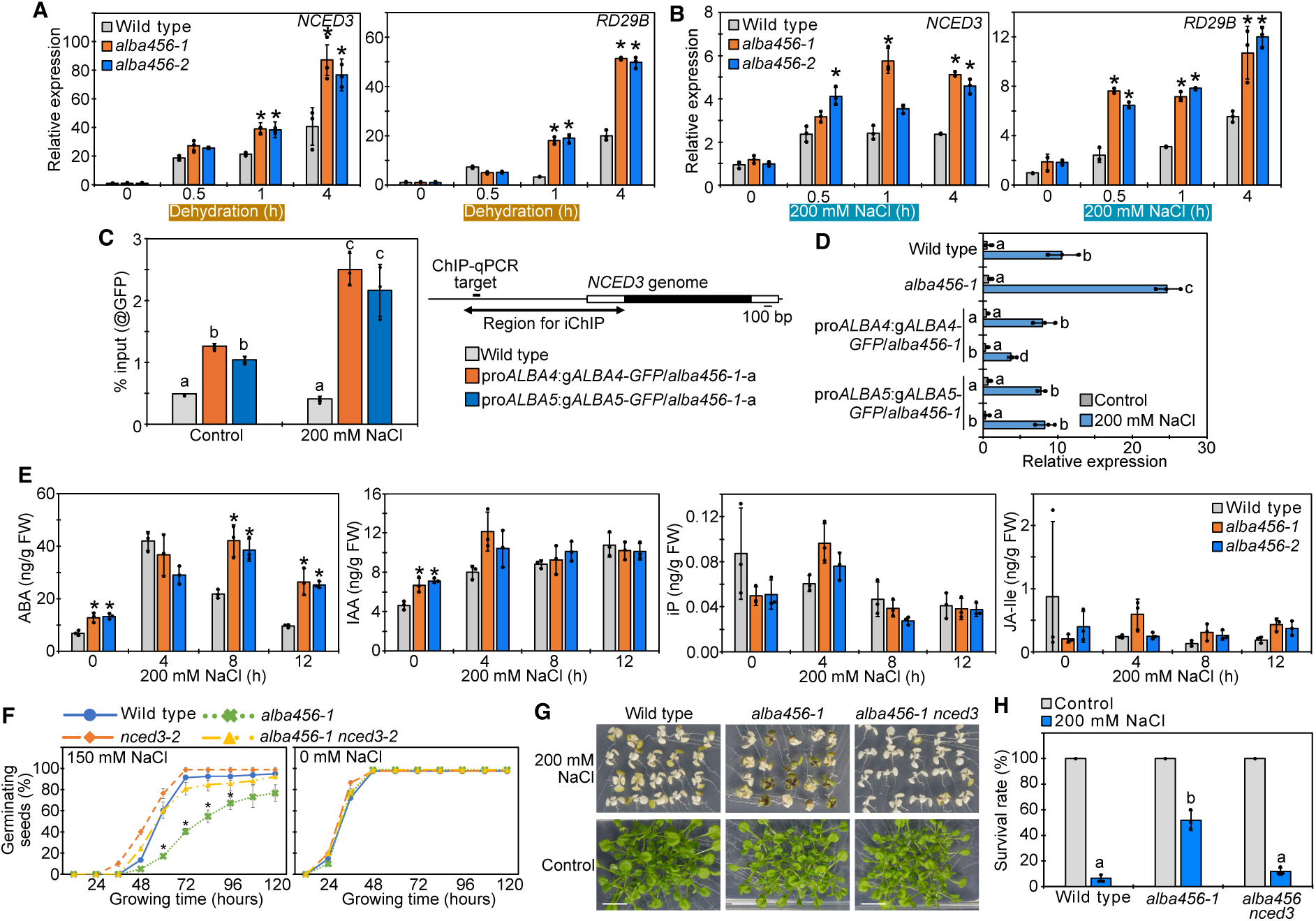
Analysis of ALBAs as novel transcriptional regulators of *NCED3* during osmotic stress conditions. (**A** and **B**) Expression levels of *NCED3* and *RD29B* under dehydration (A) and salt stress (B) conditions in *alba456* triple mutants. Error bars indicate SD from triplicate technical repeats. Asterisks indicates significant differences from wild type at each time point (*P* < 0.05, Tukey’s multiple range test). (**C**) ChIP assays of the ALBA4 and ALBA5 proteins. A schematic diagram displays the amplified region of the *NCED3* promoter. Error bars indicate SD from total six technical replicates from two independent biological replicates. Letters above bars indicate significant differences (*P* < 0.05, Tukey’s multiple range test). (**D**) Expression levels of *NCED3* under salt stress conditions in the complemented lines of ALBA4 and ALBA5 into the *alba456* triple mutant (pro*ALBA4*:g*ALBA4-GFP*/*alba456-1*-a, b and pro*ALBA5*:g*ALBA5-GFP*/*alba456-1*-a, b). Error bars indicate SD from triplicate technical repeats. Letters indicates significant differences (*P* < 0.05, Tukey’s multiple range test). (**E**) Plant hormone contents of the *alba456* triple mutants under salt stress conditions. Error bars indicate SD from three biological replicates. Asterisks indicates significant differences from wild type (*P* < 0.05, Tukey’s multiple range test). (**F**) Germination ratios of the *alba456 nced3* quadruple mutant under slat stress conditions. Error bars indicate SD from three biological replicates. Asterisks indicates significant differences in the *alba456-1* triple mutants from other plant lines (*P* < 0.05, Tukey’s multiple range test). (**G** and **H**) Salt stress tolerance of the *alba456 nced3* quadruple mutant. Images with and without salt stress (G) and the survival ratios (H) are shown. Error bars indicate SD from three biological replicates. Letters indicates significant differences during salt stress (*P* < 0.05, Tukey’s multiple range test).

### ALBAs directly bind to and suppress *NCED3* during osmotic stress

We examined whether ALBA4/5/6 affect gene expression of *NCED3*. The results showed that *NCED3* and a typical ABA-responsive gene *RD29B* (*26*) in *alba456* had significantly higher expression during dehydration and salt stresses compared to wild type (Fig. 2, A and B), consistent with the phenotypes of *alba456* (Fig. 1). Using the ALBA4-GFP and ALBA5-GFP complementation lines in *alba456-1* with their native promoters, direct binding of ALBA4/5 to the *NCED3* promoter region used in iChIP was confirmed by chromatin immunoprecipitation followed by quantitative PCR (ChIP-qPCR) (Fig. 2C). Importantly, ALBA binding was detected under both control and salt stress conditions, indicating that ALBAs are pre-associated with the *NCED3* promoter prior to transcriptional activation. The ALBA4 and ALBA5 complementation lines in the triple mutant resulted in rescued gene expression of *NCED3* during salt stress relative to the triple mutant (Fig. 2D). Hormone analysis further demonstrated that the *alba456* triple mutants had significantly higher accumulation of ABA during salt stress while amounts of other hormones were not significantly affected (Fig. 2E). These data suggest that ALBA4/5/6 directly bind to and suppress the *NCED3* expression during osmotic stress. To evaluate the genetic interaction between *ALBA4/5/6* and *NCED3*, a quadruple mutant of *alba456-1 nced3-2* was generated using a knockout mutant of *NCED3* (*27*). As a result, the quadruple mutant showed rescued phenotypes compared to the *alba456-1* triple mutant alone in germination assays during salt stress and a salt stress tolerance test (Fig. 2, F to H), suggesting that ALBA4/5/6 function upstream of *NCED3*. Taken together, these data indicated that ALBA4/5/6 are direct regulators of *NCED3* during osmotic stress to constrain its inducibility during osmotic stress.

### ALBAs show osmotic-stress-inducible nuclear condensates

To test how ALBA proteins respond to osmotic stress conditions, subcellular localization of ALBA4 was observed in the complemented pro*ALBA4*:g*ALBA4-GFP*/*alba456-1* plants crossed with a transgenic plant expressing a nuclear marker (pro*RPS5a*:*H2B-mRuby*). As previously reported, ALBA4-GFP was mainly localized in the cytoplasm under control conditions (*24*), however, we found that the salt stress conditions induced speckles of ALBA4-GFP in both nuclei and cytoplasm (Fig. 3, A to C). The speckles of ALBA-GFP were dispersed by removing salt stress conditions, and the formation of speckles was also induced by polyethylene glycol (PEG) and mannitol treatment, indicating responsiveness to osmotic stress rather than ionic toxicity (Fig. 3, A and B). ALBA5-GFP also showed formation of nuclear condensates under salt stress conditions and dispersed by removing stress conditions (fig. S3A), while the number of nuclear speckles of ALBA5-GFP was relatively lower than ALBA4-GFP (Fig. 3B and fig. S3B), suggesting functional diversification among ALBA family members. As a previous study showed ALBA5 formed condensates through liquid-liquid phase separation (LLPS) under heat stress conditions for the regulation of mRNA stability (*24*), heat stress induced ALBA5-GFP condensates in both cytoplasm and nuclei (fig. S3, A and B). ALBA4-GFP did not form condensates under heat and cold stress conditions (fig. S3C), suggesting that ALBA4 and ALBA5 have abiotic stress specificity in the formation of nuclear speckles. To assess how rapidly ALBA4-GFP forms the condensates in response to the salt stress, time-lapse imaging was performed. The result indicated that ALBA4-GFP condensates started to be detected in both nuclei and cytoplasm a few minutes after the stress conditions was started (Fig. 3D and Video S1). Next, we tested whether osmotic-stress inducible ALBA4-GFP condensates in nuclei were formed through LLPS. First, fluorescence recovery after photobleaching (FRAP) experiments showed that nuclear ALBA4-GFP signals in bleached areas recovered over time (Fig. 3, E and F, and Video S2). Next, ALBA4-GFP nuclear speckles showed dynamics and fused to each other when they came into close proximity (Fig. 3G and Video S3). Additionally, 1,6-hexandiol, which disrupts liquid-like droplets (*28*), inhibited the formation of nuclear condensation of ALBA4-GFP during salt stress (fig. S3D). These data indicated that the nuclear speckles of ALBA4-GFP in nuclei in response to salt stress were formed through LLPS. To further examine whether formation of ALBA4 condensates is involved in osmotic stress perception in plants, ALBA4-GFP was expressed in *Escherichia coli* (*E. coli*) and the ALBA4-GFP recombinant proteins were purified. The formation of ALBA4-GFP condensates in *E. coli* cells under salt stress conditions suggested that the ALBA4-GFP condensates may be formed independent of other proteins (fig. S3E). Water-deficit stress conditions, including environmental salt and dehydration stresses, are known to induce molecular crowding, where *in vivo* cell volume shrinkage induces the increase of intracellular molecular concentrations (*20*). *In vitro*, macromolecules such as PEG function as crowding agents to induce LLPS by restricting conformational space (*20*). The purified ALBA4-GFP proteins showed spherical droplets *in vitro* with a crowding agent PEG6000 (Fig. 3, H and I), suggesting that the ALBA4 proteins are sensitive to crowding independent of other proteins. Consistent with this notion, ALBA4-GFP proteins showed larger droplets as the protein concentration was increased (Fig. 3, J and K). FRAP experiments also confirmed that ALBA4-GFP signals in bleached areas recovered over time *in vitro* (fig. S3, F and G). Taken together, these data suggest that ALBA4 can function to perceive osmotic stress associated with molecular crowding in cells through LLPS.

**Fig. 3.**
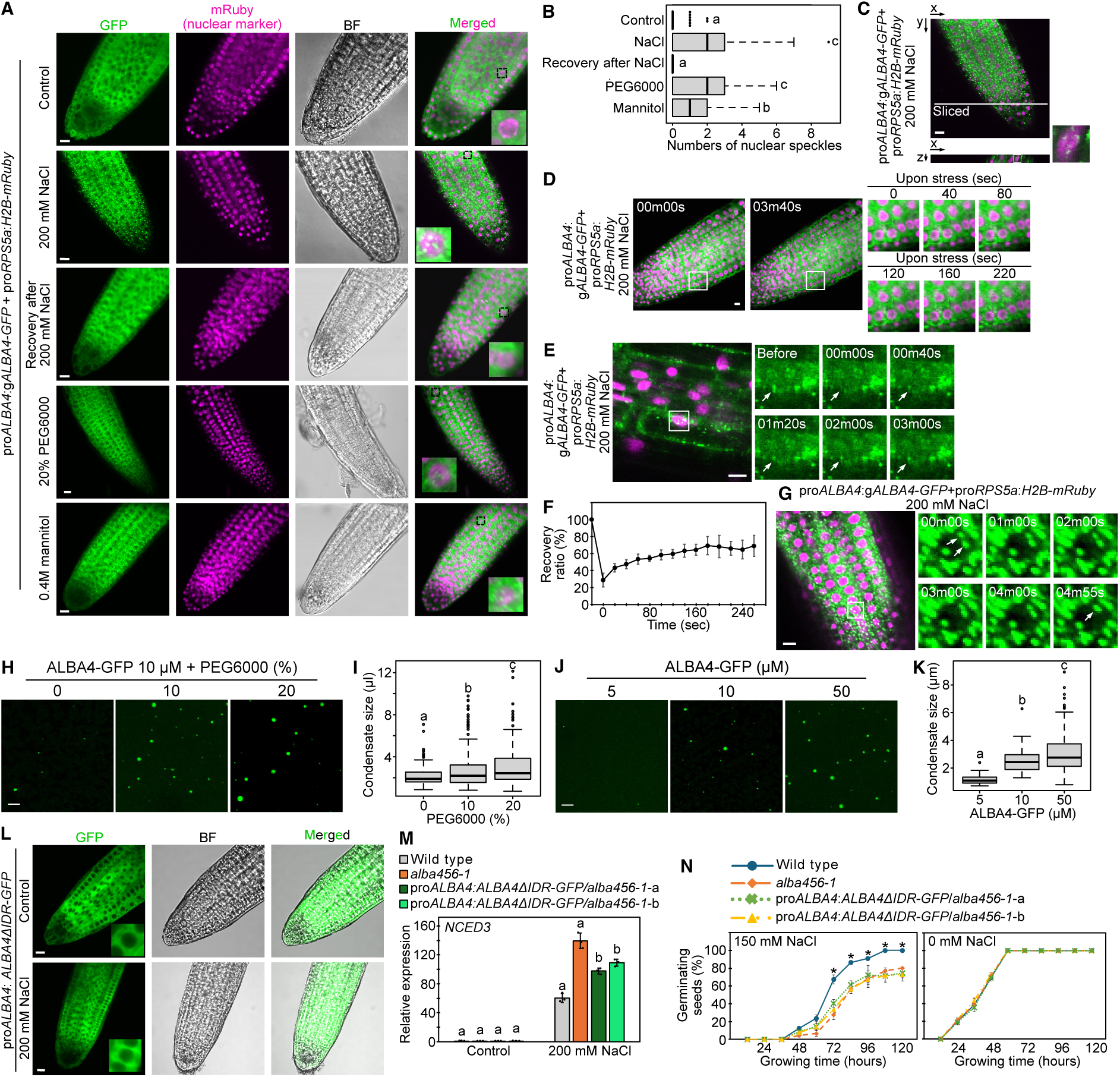
Analysis of formation of ALBA4 condensates in response to osmotic stress. **(A and B)** Confocal microscopic analysis of ALBA4-GFP under stress conditions. Confocal images (A) and numbers of nuclear speckles under each condition (B) are shown. H2B-mRuby was expressed as a nuclear marker. Scale bars represent 10 μm. Letters indicates significant differences (*P* < 0.05, Tukey’s multiple range test). (**C**) Nuclear localization of the ALBA4 speckles during salt stress. The orthogonal view of the XZ plane is shown (Scale bar: 10 μm). (D) Dynamics of formation of the ALBA4 condensates upon salt stress (Scale bar: 10 μm). (**E** and **F**) FRAP of ALBA4 nuclear condensates during salt stress *in planta*. Time-lapse images (E) and plot showing the recovery after photobleaching (F) are shown. The white arrow indicates a bleached condensate (Scale bar: 10 μm). Error bars indicate SD from twelve independent experiments. (**G**) Fusion of ALBA4 condensates in a nucleus during salt stress. The white arrows indicate condensates before/after fusion (Scale bar: 10 μm). (**H** to **K**) *In vitro* phase separation assay of ALBA4-GFP according to the presence of PEG6000 (H and I) and protein concentration (J and K). Confocal images (H and J) and the condensate size (I and K) are shown. Scale bars represent 5 μm. (**L**) Confocal microscopic analysis of root tip cells expressing ALBA4 lacking IDR under control and stress conditions (Scale bars: 10 μm). (**M**) Expression levels of *NCED3* under salt stress conditions in the complemented lines of ALBA4 lacking IDR into the *alba456* triple mutant (pro*ALBA4*:g*ALBA4ΔIDR-GFP*/*alba456-1*-a, b). Error bars indicate SD from triplicate technical repeats. Letters indicates significant differences (*P* < 0.05, Tukey’s multiple range test). (**N**) Germination ratios in the pro*ALBA4*:g*ALBA4ΔIDR-GFP*/*alba456-1* plants under slat stress conditions. Error bars indicate SD from three biological replicates. Asterisks indicates significant differences in wild type from other lines (*P* < 0.05, Tukey’s multiple range test).

### ALBA4 IDR is necessary for osmotic stress responses in plants

It is predicted that ALBA4 has an intrinsically disordered region (IDR) at the C terminus (*24*), and it is generally known that IDRs can be necessary for the formation of condensates (*20*). To test whether the IDR of ALBA4 is also required for formation of condensates in response to osmotic stress, full-length or truncated ALBA4 proteins fused to GFP were expressed in *Nicotiana benthamiana* (tobacco) leaves (fig. S3H). Even in the tobacco cells, the full-length ALBA4-GFP protein formed nuclear speckles under salt stress conditions (fig. S3, I and J). Deletion of the C-terminal IDR (ALBA4ΔIDR) significantly suppressed the formation of nuclear condensates while deletion of the N-terminal Alba domain (ALBA4ΔN) still showed nuclear speckles (fig. S3, I and J), suggesting that the C-terminal IDR of ALBA4 is sufficient to form condensates in response to osmotic stress. On the basis of this result, the ALBA4ΔIDR was introduced into the *alba456-1* triple mutant under the control of the native promoter (pro*ALBA4*:*ALBA4ΔIDR-GFP*/*alba456-1*). ALBA4ΔIDR-GFP did not show any condensates in both nucleus and cytoplasm in Arabidopsis (Fig. 3L). Additionally, the triple mutant complemented with ALBA4ΔIDR did not fully rescue the gene expression of *NCED3* and the delayed germination phenotypes of *alba456-1* under salt stress conditions (Fig. 3, M and N). These data suggested that the IDR of ALBA4 is necessary for LLPS and for full functionality of ALBA4 to suppress the *NCED3* gene expression under salt stress conditions.

As previous studies indicated that ALBAs regulate RNA stability in cytoplasm (*24, 29*), we evaluated how the cytosolic ALBA4 condensates during osmotic stress affect the *NCED3* mRNA stability and phenotypes. First, ALBA4-GFP was co-expressed in tobacco leaves under salt stress conditions with DCP1-RFP or RBP47B-RFP, which are markers of processing bodies (PBs) or stress granules (SGs), respectively (*30, 31*). ALBA4 condensates in the cytoplasm showed almost complete colocalization with RBP47B while they were partially colocalized with DPC1 (fig. S4, A to C), suggesting that the cytosolic ALBA4 proteins were mainly localized in SGs as reported previously (*24, 29*). Additionally, we found that the *NCED3* mRNA during salt stress showed a higher rate of degradation in the *alba456* triple mutant relative to wild type (fig. S4D) similar to reported functions of cytosolic ALBA4/5/6 toward the target mRNAs during heat stress (*24*), suggesting that the suppressive effects of ALBA4/5/6 on the gene expression of *NCED3* should be mainly present in nuclei. To test this notion, we complemented the *alba456-1* triple mutant with ALBA4-GFP fused to nuclear localization signal (nls) (pro*ALBA4*:g*ALBA4-GFP-nls*/*alba456-1*). The ALBA4-GFP-nls complemented lines significantly rescued phenotypes of the *alba456-1* triple mutant (fig. S4, E to G). These data suggest that nuclear-localized ALBA4 has a significant contribution to plant phenotypes under salt stress conditions.

### ALBAs suppress transcriptomic changes during osmotic stress

To reveal how ALBA4/5/6 proteins affect transcriptomics under osmotic stress conditions, we performed RNA sequencing (RNA-seq) in *alba456-1* and wild-type under control, dehydration, and salt stress conditions (tables S5, S6 and S7). Larger numbers of the upregulated genes in *alba456-1* relative to the downregulated genes especially during the stress conditions implied predominant suppressive effects of ALBAs on stress-induced transcriptional responses, consistent with the effects on *NCED3* and *RD29B* (Fig. 4A and Fig. 2, A and B). Next, to evaluate how the *NCED3*-dependent pathway was affected in *alba456-1*, we compared the published transcriptomic data in the *NCED3* knockout mutant (*nc3-2*) at multiple timepoints during dehydration stress (*27*). The results showed that downregulated genes in *nc3-2* were significantly enriched among upregulated genes in *alba456-1* during the dehydration and salt stress (Fig. 4, B to D), while relatively smaller overlaps were detected between the upregulated genes in *nc3-2* and downregulated genes in *alba456-1* (Fig. 4E), consistent with the upregulation of *NCED3* in *alba456-1* (Fig. 2, A and B). Additionally, the upregulated genes in *alba456-1* during the dehydration and salt stress showed significantly large overlaps with dehydration- and salt-stress-inducible genes in wild-type, respectively (Fig. 4, F and G) compared to the overlaps between the stress-suppressive genes and downregulated genes in *alba456-1* during the stress conditions (fig. S5, A and B, tables S8 and S9). These results suggest that ALBA4/5/6 mainly has suppressive effects on stress-dependent transcriptomic changes, especially on stress-activated genes partially through suppression of *NCED3*. Gene ontology (GO) analysis also showed that ABA- and osmotic-stress-related ontologies were significantly enriched among the upregulated genes in *alba456-1* during both dehydration and salt stresses (Fig. 4, H and I) while cytokinin- and auxin-related ontologies (fig. S5C) and dark condition-related ontology (fig. S5D) were enriched among the downregulated genes during the dehydration and salt stress, respectively, consistent with the higher stress tolerance and inhibited growth under the stress conditions in *alba456-1* (Fig. 1). The upregulated and downregulated genes in *alba456-1* under control conditions were enriched with iron homeostasis- and circadian-related ontologies (fig. S5E) and nitrate-related ontologies (fig. S5F), respectively, suggesting additional functions of ALBA4/5/6 under non-stress conditions. These may also be correlated with the circadian-dependent phenotypes of leaf temperature in *alba456-1* (Fig. 1, L and M). *Cis*-element enrichment analysis showed that abscisic acid-responsive elements (ABRE) (ACGTGG/TC)-related motifs and the core G box (CACGTG) which were bound by typical ABA-responsive transcription factors ABSCISIC ACID-RESPONSIVE ELEMENT-BINDING PROTEINs/ABRE-BINDING FACTORs (AREBs/ABFs) (*32*), were highly enriched among the 1-kb promoter regions of the upregulated genes in *alba456-1* during the dehydration and salt stress (Fig. 4, J and K), while TATA-related motifs were enriched among the downregulated genes in *alba456-1* (fig. S5, G and H). Taken together, these data indicate that ALBA4/5/6 functions to suppress stress-inducible genes, including *NCED3*, during osmotic stress.

**Fig. 4.**
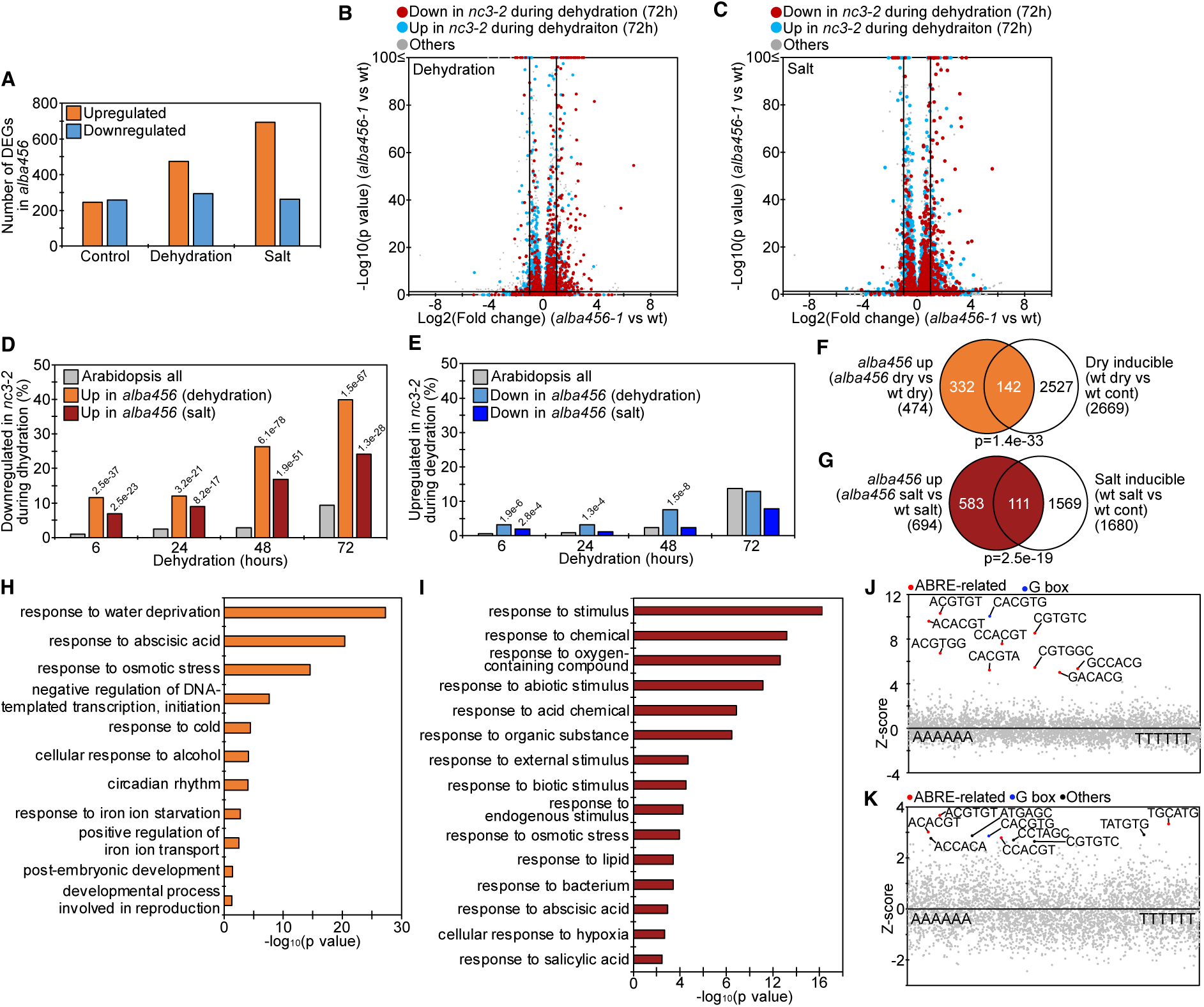
Transcriptomic analysis of *alba456-1* triple mutant under osmotic stress conditions. (**A**) Number of differentially expressed genes in *alba456-1* under control, dehydration and salt stress conditions relative to wild type (|Log_2_FC| ≥ 1, FDR < 0.05). (**B** and **C**) Volcano plots showing DEGs in *alba456-1* during dehydration (B) and salt (C) stress with downstream genes of *NCED3*. Red and blue dots represent down- and upregulated genes in the *nced3* knockout mutant (*nc3-2*), respectively, according to the published paper (*27*). Vertical and horizontal lines mark thresholds of up- and downregulated genes in *alba456-1* are shown (|Log_2_FC| ≥ 1, FDR <0.05). (**D** and **E**) Plots show the percentage of up- (D) and downregulated (E) genes in *nc3-2* at each time point of dehydration stress (*27*) among Arabidopsis all genes and DEGs in *alba456-1* during dehydration and salt stress. Values on the bars represent p values from Arabidopsis all (pairwise Fisher’s exact test with Benjamini–Hochberg correction). (**F** and **G**) Venn diagrams showing overlaps between stress inducible genes and upregulated genes in *alba456-1* during dehydration (F) and salt (G) stress. P values are calculated by Fisher’s exact test. (**H** and **I**) Enrichment of GOs among upregulated genes in *alba456-1* during dehydration (H) and salt (I) stress. (**J** and **K**) Overrepresentation analysis of hexamer motifs in the 1-kb promoter regions of the top 200 upregulated genes in *alba456-1* during dehydration (J) and salt (K) stress. Z scores (y axis) for the observed frequencies of all hexamer motifs (x axis) are presented in the scatter plot. The ABRE-related, G box and other top 10 enriched motifs are highlighted, in red, blue and black, respectively.

### ALBA4 restricts osmotic stress inducibility of the target genes

To reveal how ALBA4 affects the direct target genes under stress conditions, we next performed ChIP-seq of ALBA4 using the pro*ALBA4*:g*ALBA4-GFP*/*alba456-1* plants under control and salt stress conditions with relatively high reproducibility of two replicates (fig. S6A and tables S10, S11 and S12), and identified ALBA4 peaks under control and salt stress conditions (Fig. 5A). There were more and higher peaks during salt stress conditions compared to control, consistent with the ChIP-qPCR (Fig. 2C) and formation of salt-inducible nuclear speckles (Fig. 3). We also found that there was a large overlap between the peaks under control and salt stress conditions (Fig. 5, A and B). We next divided the peaks into three groups: control-specific, salt-specific, and common peaks (Fig. 5B). All of the three groups of peaks were mainly located around promoter regions (Fig. 5C), suggesting functions of ALBA4 as a transcriptional regulator. Additionally, the three groups of peaks were enriched with the G box motif (CACGTG) or GGGCCC elements (Fig. 5D). As previous studies suggested that ALBA proteins do not have DNA-binding activity to specific motifs (*23*), these results suggest that ALBA4 is recruited to target loci through interactions with transcription factors bound to these motifs. Next, we identified direct target genes associated with the ALBA4 peaks (Fig. 5, E and F). Similarly, the ALBA4-bound genes under the control and salt stress conditions showed a large overlap, and we divided them into three groups: control-specific, salt-specific and common ALBA4-bound genes (Fig. 5F). Importantly, the common ALBA4-bound genes were enriched with GOs related to stress responses similar to the salt-specific ALBA4-bound genes while the control-specific ALBA4-bound genes were enriched with GOs related to development (fig. S6, B to D), suggesting that ALBA4 is located to stress-related genes even before the onset of stress conditions. This is consistent with the ChIP-qPCR result showing that ALBA4 bound to the *NCED3* promoter under both control and salt stress conditions (Fig. 2C). Next, we identified direct target genes of ALBA4 among DEGs in *alba456-1* under control, salt, and dehydration stress conditions (Fig. 5G). We also examined the DEGs in *alba456-1* during dehydration stress, considering that high salt and dehydration stress are known to induce common ABA-dependent molecular mechanisms due to the high osmotic conditions (*33, 34*). The result showed that ALBA4 directly influenced more genes under the stress condition relative to the control condition mainly through suppression of target genes compared to activation (Fig. 5G). Both salt-specific and common ALBA4-bound genes among upregulated genes under the salt stress condition in *alba456-1* were enriched with GOs related to water deprivation (Fig. 5, H and I) while the ALBA4 directly suppressed genes under the control condition were enriched with GOs related to circadian rhythm and iron homeostasis (fig. S6E). The 1-kb promoter regions of the ALBA4 directly suppressed genes during salt stress and potentially during dehydration stress were enriched with G box-related motifs while no enriched motifs were detected within the 1-kb downstream regions (fig. S6, F and G). Next, we examined whether the ALBA4 directly suppressed genes were induced or suppressed under the stress conditions. As a result, we found that salt-stress-inducible genes were enriched within the directly suppressed genes compared to all upregulated genes during the salt stress while salt-stress-suppressive genes were not enriched (Fig. 5J). Additionally, we found that the ALBA4-bound genes showed significantly lower stress inducibility during the salt stress among all salt-inducible genes in Arabidopsis (Fig. 5K), suggesting that ALBA4 may be a determinant of gene inducibility in response to salt stress. The same tendency was also confirmed within the potentially ALBA4 directly suppressed genes during dehydration stress: dehydration-inducible genes were enriched and the potential binding of ALBA4 was associated with significantly lower dehydration stress inducibility (fig. S6, H and I). Taken together, these data suggest that ALBA4 has direct suppressive effects on salt-stress-inducible genes during salt stress on a genome-wide level, binding to them even under control conditions. This model is consistent with the functions of ALBAs on the *NCED3* gene (Fig. 2). Importantly, the list of ALBA4-direct suppressive genes includes not only *NCED3* (Fig. 5L) but also other well studied drought- and salt-stress-inducible genes such as *Galactinol Synthase 2* (*GolS2*) (*35*) (Fig. 5M), *ABF3* (*32*), *MITOGEN-ACTIVATED PROTEIN KINASE KINASE KINASE 17* (*MAPKKK17*) and *MAPKKK18* (*36*) (fig. S6, J to L), suggesting that ALBA4 has critical roles in response to salt stress through regulation of these genes as a transcriptional regulator.

**Fig. 5.**
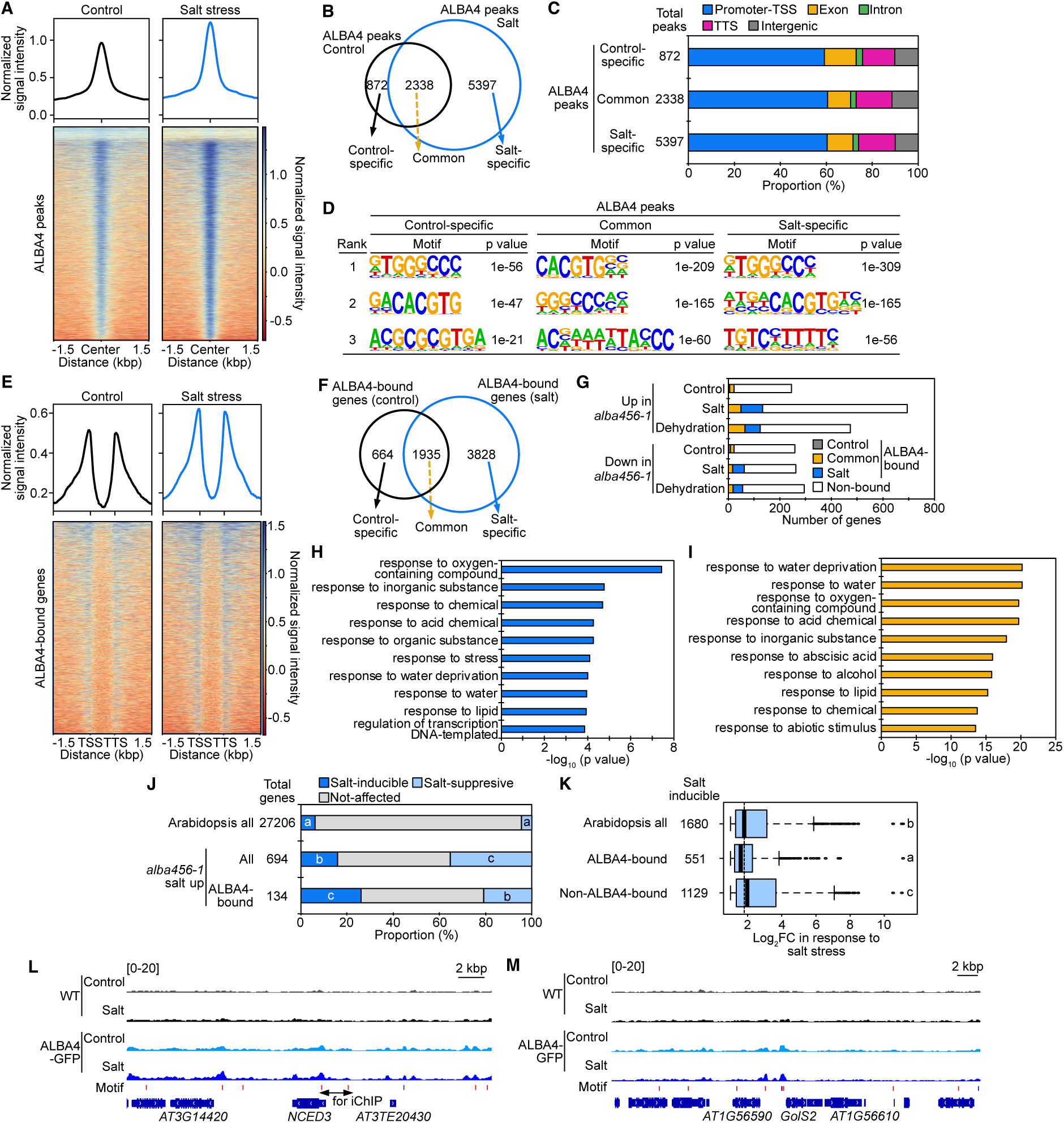
ALBA4 suppresses osmotic stress inducibility of the target genes. (**A**) Metagene plots and heatmaps showing the ALBA4 ChIP-seq peaks and flanking regions under control and salt stress conditions using the pro*ALBA4*:g*ALBA4-GFP*/*alba456-1* plant. (**B**) Venn diagrams showing overlaps of ALBA4 ChIP-seq peaks between control and salt stress. (**C**) Plots showing the proportions of ALBA4 peaks at annotated genic and intergenic regions. (**D**) Enriched motifs identified by MEME-CHIP using the three groups of ALBA4 ChIP-seq peaks in Fig. 5B. (**E**) Metagene plots and heatmaps showing the ALBA4 ChIP-seq peaks around target genes and flanking regions under control and salt stress conditions using the pro*ALBA4*:g*ALBA4-GFP*/*alba456-1* plant. TSS and TTS represent transcription start site and transcription termination site, respectively. (**F**) Venn diagrams showing overlaps of ALBA4-bound genes between control and salt stress. (**G**) Plot showing the numbers of DEGs in *alba456-1* under control, salt and dehydration stress conditions which bound by ALBA4 under control and salt stress conditions. (**H** and **I**) Enrichment of GOs among upregulated genes in *alba456-1* during salt stress which were bound by ALBA4 under salt-specific (H) and both control and salt (I) conditions. (**J**) Plots showing proportions of stress inducible genes among Arabidopsis all genes and upregulated genes in *alba456-1* during salt stress which were bound by ALBA4. Letters represent significant differences within the proportions of stress-inducible or suppressive genes (*P* < 0.05, pairwise Fisher’s exact test with Benjamini–Hochberg correction). (**K**) Plots showing salt stress inducibility of genes bound by ALBA4. Letters represent significant differences (*P* < 0.05, pairwise Wilcoxon test with Benjamini–Hochberg correction). (**L** and **M**) IGV screenshots showing the ChIP-seq signals of ALBA4 on *NCED3* (L) and *GolS2* (M). All tracks represent read density normalized to sequencing depth. The gene models are shown at the bottom with arrow heads showing the direction of transcription. Red and blue bars represent locations of G box (CACGTG) and GGGCCC motifs, respectively.

## Discussion

Molecular mechanisms of osmotic stress responses in plants at early timepoints are still largely unknown. In this study, using the iChIP screen method *in planta*, we identified ALBA proteins and revealed that formation of ALBA condensates in nuclei and cytoplasm could function as a potential osmo-sensing mechanism with molecular crowding to regulate ABA biosynthesis. So far, potential osmo-sensing mechanisms such as changes of lipid membrane fluidity and protein structure, or oxidative signaling from mitochondria and chloroplasts have been suggested in plants (*1, 6, 37*). Recent studies have shown that sensing molecular crowding through LLPS is another type of osmo-sensing mechanism during cell shrinkage under osmotic conditions. In Arabidopsis, the transcription factor SEU (*18*) and mRNA decapping enzyme DCP5 (*19*) were revealed to function as osmo-sensors through LLPS, while, in human cells, apoptosis signal-regulating kinase 3 (ASK3) was identified (*38*). Our study reveals how osmotic-inducible LLPS is linked to ABA biosynthesis in the early stress response through ALBA proteins especially through restricting gene inducibility. ALBA4 showed formation of condensates through LLPS both *in vivo* and *in vitro* upon salt stress (Fig. 3). Additionally, consistent with IDR requirement for ALBA5 to form condensates in SGs under heat stresses (*24*), the IDR of ALBA4 was necessary to form condensates in response to osmotic stresses (Fig. 3). A phylogenetic analysis revealed that the RGG enriched IDRs at the C terminal region of ALBA4 are conserved across kingdoms (*23*), suggesting that common osmo-sensing mechanisms with RGG-enriched IDRs might exist in various organisms.

Nuclear compartmentalization through LLPS is involved in various molecular mechanisms, including transcription, epigenetic regulation of histone modifications and chromatin conformation in various organisms, and, in plants, stress-inducible nuclear speckles have been reported during heat, cold, and osmotic stresses (*39, 40*). In all of these processes, specific regulatory factors are selectively recruited to or excluded from the compartmentalized nuclear regions (*41, 42*). Considering that ALBA4 nuclear speckles mainly showed negative effects on target gene expression (Fig. 2, Fig. 4, and Fig. 5), any active factors and/or negative factors may be excluded and/or recruited to the ALBA condensates, respectively. The osmotic-inducible ALBA condensates are also located in both nuclei and SG in cytoplasm (fig. S4). It is suggestive that components of SG in mammals have been reported to shuttle between cytoplasm and nucleus (*43*), and some of them, including the Fused in sarcoma (FUS) protein, involve transcriptional regulation in nuclear speckles through LLPS (*44*). In plants, DCP5 interacts with ALBA proteins, is located in PBs (*24*), and it is also reported to function as a transcriptional suppressor in nuclear speckles through LLPS during flowering (*45*). As a recently published paper in rice reports that SGs including the DROUGHT RESISTANCE GENE 9 (DRG9) protein protect *OsNCED4* mRNA from degradation (*46*), interestingly, our data also indicated that ALBA proteins increased the stability of the *NCED3* mRNA during stress conditions (fig. S4D). This suggests evolutionarily conserved functions of SG components across plants and mammals, which involve both transcription in nuclei and post-transcriptional regulation of RNA in the cytoplasm, while how the mechanistic linkage between these two processes remains unclear.

In this study, we applied the iChIP screen method (*21*) to isolate transcriptional regulators at a specific promoter *in planta* (Fig. 1). This can be a powerful tool to reveal transcriptional mechanisms at a specific promoter, similar to other corresponding methods of proteomics on a specific genome locus (*47*). Especially, the promoter regions of direct ABLA4 target genes were significantly enriched with G boxes (fig. S6F), suggesting G box-binding transcription factors, which work together with ALBAs. This is potentially related to transcriptional regulation of *NCED3*, whose promoter also contains a G box motif required for the gene activation (*22*). Future studies are needed to analyze other candidate transcriptional regulators identified in our iChIP screen (table S1) to elucidate the detailed molecular mechanisms of ABA biosynthesis through ALBAs.

Overall, our findings provide a molecular model of the early response to osmotic stress in plants before ABA accumulation. ALBA4/5/6, potential osmo-sensing proteins through molecular crowding, suppress inducibility of target osmotic stress-inducible genes to balance between stress tolerance and plant growth, especially through suppression of *NCED3* (fig. S6M). Our findings will facilitate various researches in the broad field of stress responses in plants.

## ACKNOWLEDGMENTS

We thank S. Mibu (The University of Tokyo, Chiba, Japan) and K. Kuwata (The Nagoya University, Nagoya, Japan) for substantial technical assistance.

## Funding

Japan Society for the Promotion of Science (MXT/JSPS KAKENHI) (Grant-in-Aid for Early-Career Scientist) 22K15137 (HS)

Japan Society for the Promotion of Science (MXT/JSPS KAKENHI) 24K09498 (HS)

Japan Society for the Promotion of Science (MXT/JSPS KAKENHI) JP22H04925 (PAGS) (HS)

Human Frontier Scientific Program fellowship grant LT000162/2018-L (HS)

Japan Society for the Promotion of Science (MXT/JSPS KAKENHI) 20H03297 (SM)

Japan Society for the Promotion of Science (MXT/JSPS KAKENHI) 22H00415 (SM)

ASPIRE JPMJAP2306 (SM)

Yakult Pharmaceutical Industry Co., Ltd

ACRO Research grant of Teikyo university TeTe20-01 (MA)

## Author contributions

H.S. conceived and designed the experiments, performed the experiments and analyzed the data. S.F. designed and performed the iChIP screen. M.S. and M.F. performed the RIPPS phenotyping analysis. D.S. performed the data analysis in the RNA-seq. E.M.S. contributed to the mass spectrometry. E.Y. and M.A. performed the hormonome analysis. A.K. and Y.S. performed the data analysis in the ChIP-seq. H.S. wrote the manuscript, and all authors revised, commented and agreed on the manuscript before submission. F.T., K.Y.-S, K.S and S.M. supervised the project.

## Data, code, and materials availability

All data are available in the manuscript, supplementary materials. All newly generated materials (plasmids, seeds) are available from the corresponding author upon reasonable request. Raw data of mass spectrometry has been deposited at jPOST Repository Database with accession number PXD075188. RNA-seq and ChIP-seq raw read data have been deposited at National Center for Biotechnology Information Gene Expression Omnibus database with accession numbers PRJDB16910 and PRJDB20689. Accession numbers of all gene sequences are listed in table S14. Previously published RNA-seq read was used in this study (*27*).

## SUPPLEMENTARY MATERIALS

Detailed Materials and Methods and Figs. S1 to S6 are shown in the Supplementary file.

## References and Notes

1. H. Sato, J. Mizoi, K. Shinozaki, K. Yamaguchi-Shinozaki, Complex plant responses to drought and heat stress under climate change. The Plant Journal 117, 1873–1892 (2024).

2. T. Yoshida, A. R. Fernie, Hormonal regulation of plant primary metabolism under drought. J Exp Bot 75, 1714–1725 (2024).

3. M. M. Chaves, J. Flexas, C. Pinheiro, Photosynthesis under drought and salt stress: regulation mechanisms from whole plant to cell. Ann Bot 103, 551–560 (2009).

4. S. I. Zandalinas, R. Mittler, D. Balfagón, V. Arbona, A. Gómez-Cadenas, Plant adaptations to the combination of drought and high temperatures. Physiol Plant 162, 2–12 (2018).

5. R. C. Nongpiur, S. L. Singla-Pareek, A. Pareek, The quest for osmosensors in plants. J Exp Bot 71, 595–607 (2020).

6. B. Yu, D. Y. Chao, Y. Zhao, How plants sense and respond to osmotic stress. J Integr Plant Biol 66, 394–423 (2024).

7. T. Kuromori, M. Seo, K. Shinozaki, ABA Transport and Plant Water Stress Responses. Trends Plant Sci 23, 513–522 (2018).

8. Y. Ma et al., Molecular Mechanism for the Regulation of ABA Homeostasis During Plant Development and Stress Responses. Int J Mol Sci 19, (2018).

9. A. Daszkowska-Golec, ABA is important not only under stress - revealed by the discovery of new ABA transporters. Trends Plant Sci 27, 423–425 (2022).

10. J. Fidler et al., PYR/PYL/RCAR Receptors Play a Vital Role in the Abscisic-Acid-Dependent Responses of Plants to External or Internal Stimuli. Cells 11, (2022).

11. F. Soma, F. Takahashi, K. Yamaguchi-Shinozaki, K. Shinozaki, Cellular Phosphorylation Signaling and Gene Expression in Drought Stress Responses: ABA-Dependent and ABA-Independent Regulatory Systems. Plants (Basel) 10, (2021).

12. S. Iuchi et al., Regulation of drought tolerance by gene manipulation of 9-cis-epoxycarotenoid dioxygenase, a key enzyme in abscisic acid biosynthesis in Arabidopsis. Plant J 27, 325–333 (2001).

13. H. Sato et al., Arabidopsis thaliana NGATHA1 transcription factor induces ABA biosynthesis by activating NCED3 gene during dehydration stress. Proc Natl Acad Sci U S A 115, E11178–E11187 (2018).

14. M. K. Jensen et al., ATAF1 transcription factor directly regulates abscisic acid biosynthetic gene NCED3 in Arabidopsis thaliana. FEBS Open Bio 3, 321–327 (2013).

15. Y. Ding, Z. Avramova, M. Fromm, The Arabidopsis trithorax-like factor ATX1 functions in dehydration stress responses via ABA-dependent and ABA-independent pathways. Plant J 66, 735–744 (2011).

16. Q. Liu, W. Liu, Y. Niu, T. Wang, J. Dong, Liquid-liquid phase separation in plants: Advances and perspectives from model species to crops. Plant Commun 5, 100663 (2024).

17. L. Shen, Epitranscriptomic regulation through phase separation in plants. Trends Plant Sci 30, 629–641 (2025).

18. B. Wang et al., Condensation of SEUSS promotes hyperosmotic stress tolerance in Arabidopsis. Nat Chem Biol 18, 1361–1369 (2022).

19. Z. Wang et al., A cytoplasmic osmosensing mechanism mediated by molecular crowding-sensitive DCP5. Science 386, eadk9067 (2024).

20. C. Alfano et al., Molecular Crowding: The History and Development of a Scientific Paradigm. Chem Rev 124, 3186–3219 (2024).

21. T. Fujita, H. Fujii, Efficient isolation of specific genomic regions retaining molecular interactions by the iChIP system using recombinant exogenous DNA-binding proteins. BMC Mol Biol 15, 26 (2014).

22. B. Behnam et al., Characterization of the promoter region of an Arabidopsis gene for 9-cis-epoxycarotenoid dioxygenase involved in dehydration-inducible transcription. DNA Res 20, 315–324 (2013).

23. M. Goyal, C. Banerjee, S. Nag, U. Bandyopadhyay, The Alba protein family: Structure and function. Biochim Biophys Acta 1864, 570–583 (2016).

24. J. Tong et al., ALBA proteins confer thermotolerance through stabilizing HSF messenger RNAs in cytoplasmic granules. Nat Plants 8, 778–791 (2022).

25. M. Fujita, T. Tanabata, K. Urano, S. Kikuchi, K. Shinozaki, RIPPS: A Plant Phenotyping System for Quantitative Evaluation of Growth Under Controlled Environmental Stress Conditions. Plant Cell Physiol 59, 2030–2038 (2018).

26. Y. Uno et al., Arabidopsis basic leucine zipper transcription factors involved in an abscisic acid-dependent signal transduction pathway under drought and high-salinity conditions. Proc Natl Acad Sci U S A 97, 11632–11637 (2000).

27. K. Urano et al., Analysis of plant hormone profiles in response to moderate dehydration stress. Plant J 90, 17–36 (2017).

28. S. Kroschwald et al., Promiscuous interactions and protein disaggregases determine the material state of stress-inducible RNP granules. Elife 4, e06807 (2015).

29. H. Ge et al., Debranching enzyme DBR1-mediated lariat RNA turnover requires ALBA proteins in Arabidopsis. Mol Cell 85, 4183–4197.e4186 (2025).

30. G. J. Jang, J. C. Jang, S. H. Wu, Dynamics and Functions of Stress Granules and Processing Bodies in Plants. Plants (Basel) 9, (2020).

31. K. Motomura et al., Diffuse decapping enzyme DCP2 accumulates in DCP1 foci under heat stress in Arabidopsis thaliana. Plant Cell Physiol 56, 107–115 (2015).

32. T. Yoshida et al., Four Arabidopsis AREB/ABF transcription factors function predominantly in gene expression downstream of SnRK2 kinases in abscisic acid signalling in response to osmotic stress. Plant Cell Environ 38, 35–49 (2015).

33. Q. Hussain et al., Transcription Factors Interact with ABA through Gene Expression and Signaling Pathways to Mitigate Drought and Salinity Stress. Biomolecules 11, (2021).

34. J. K. Zhu, Salt and drought stress signal transduction in plants. Annu Rev Plant Biol 53, 247–273 (2002).

35. T. Taji et al., Important roles of drought- and cold-inducible genes for galactinol synthase in stress tolerance in Arabidopsis thaliana. Plant J 29, 417–426 (2002).

36. S. W. Choi, S. B. Lee, Y. J. Na, S. G. Jeung, S. Y. Kim, Arabidopsis MAP3K16 and Other Salt-Inducible MAP3Ks Regulate ABA Response Redundantly. Mol Cells 40, 230–242 (2017).

37. G. Miller, N. Suzuki, S. Ciftci-Yilmaz, R. Mittler, Reactive oxygen species homeostasis and signalling during drought and salinity stresses. Plant Cell Environ 33, 453–467 (2010).

38. K. Watanabe et al., Cells recognize osmotic stress through liquid-liquid phase separation lubricated with poly(ADP-ribose). Nat Commun 12, 1353 (2021).

39. D. Fu, B. Jiang, Liquid-liquid phase separation regulates gene expression in plants. Agriculture Communications, 100084 (2025).

40. S. Fujishiro, M. Sasai, K. Maeshima, Chromatin domains in the cell: Phase separation and condensation. Curr Opin Struct Biol 91, 103006 (2025).

41. A. S. Belmont, Nuclear Compartments: An Incomplete Primer to Nuclear Compartments, Bodies, and Genome Organization Relative to Nuclear Architecture. Cold Spring Harb Perspect Biol 14, (2022).

42. Y. R. Kamimura, M. Kanai, Chemical insights into liquid-liquid phase separation in molecular biology. Bulletin of the Chemical Society of Japan 94, 1045–1058 (2021).

43. N. Kedersha, P. Anderson, Regulation of translation by stress granules and processing bodies. Prog Mol Biol Transl Sci 90, 155–185 (2009).

44. S. Reber et al., The phase separation-dependent FUS interactome reveals nuclear and cytoplasmic function of liquid-liquid phase separation. Nucleic Acids Res 49, 7713–7731 (2021).

45. W. Wang et al., The P-body component DECAPPING5 and the floral repressor SISTER OF FCA regulate FLOWERING LOCUS C transcription in Arabidopsis. Plant Cell 35, 3303–3324 (2023).

46. H. Wang et al., A double-stranded RNA binding protein enhances drought resistance via protein phase separation in rice. Nat Commun 15, 2514 (2024).

47. S. L. Xu, R. Shrestha, S. S. Karunadasa, P. Q. Xie, Proximity Labeling in Plants. Annu Rev Plant Biol 74, 285–312 (2023).

48. T. Murashige, F. Skoog, A revised medium for rapid growth and bio assays with tobacco tissue cultures. Physiologia plantarum 15, (1962).

49. A. Horie et al., SWI/SNF chromatin remodeling factor BRAHMA promotes de novo shoot regeneration by epigenetic priming via H3K27me3 removal. Plant J 124, e70630 (2025).

50. M. K. Shibuta et al., A live imaging system to analyze spatiotemporal dynamics of RNA polymerase II modification in Arabidopsis thaliana. Commun Biol 4, 580 (2021).

51. S. J. Clough, A. F. Bent, Floral dip: a simplified method for Agrobacterium-mediated transformation of Arabidopsis thaliana. Plant J 16, 735–743 (1998).

52. F. Fauser, S. Schiml, H. Puchta, Both CRISPR/Cas-based nucleases and nickases can be used efficiently for genome engineering in Arabidopsis thaliana. Plant J 79, 348–359 (2014).

53. R. P. Hellens, E. A. Edwards, N. R. Leyland, S. Bean, P. M. Mullineaux, pGreen: a versatile and flexible binary Ti vector for Agrobacterium-mediated plant transformation. Plant Mol Biol 42, 819–832 (2000).

54. H. Sato, J. Santos-González, C. Köhler, Combinations of maternal-specific repressive epigenetic marks in the endosperm control seed dormancy. Elife 10, (2021).

55. H. Sato et al., Arabidopsis DPB3-1, a DREB2A interactor, specifically enhances heat stress-induced gene expression by forming a heat stress-specific transcriptional complex with NF-Y subunits (vol 26, pg 4954, 2014). Plant Cell 27, 2076–2077 (2015).

56. F. Soma et al., ABA-unresponsive SnRK2 protein kinases regulate mRNA decay under osmotic stress in plants. Nat Plants 3, 16204 (2017).

57. T. N. Uehara et al., Casein kinase 1 family regulates PRR5 and TOC1 in the Arabidopsis circadian clock. Proc Natl Acad Sci U S A 116, 11528–11536 (2019).

58. A. M. Bolger, M. Lohse, B. Usadel, Trimmomatic: a flexible trimmer for Illumina sequence data. Bioinformatics 30, 2114–2120 (2014).

59. A. Dobin et al., STAR: ultrafast universal RNA-seq aligner. Bioinformatics 29, 15–21 (2013).

60. Y. Liao, G. K. Smyth, W. Shi, featureCounts: an efficient general purpose program for assigning sequence reads to genomic features. Bioinformatics 30, 923–930 (2014).

61. M. I. Love, W. Huber, S. Anders, Moderated estimation of fold change and dispersion for RNA-seq data with DESeq2. Genome Biol 15, 550 (2014).

62. S. X. Ge, D. Jung, R. Yao, ShinyGO: a graphical gene-set enrichment tool for animals and plants. Bioinformatics 36, 2628–2629 (2020).

63. K. Maruyama et al., Identification of cis-acting promoter elements in cold- and dehydration-induced transcriptional pathways in Arabidopsis, rice, and soybean. DNA Res 19, 37–49 (2012).

64. S. Oya, M. Takahashi, K. Takashima, T. Kakutani, S. Inagaki, Transcription-coupled and epigenome-encoded mechanisms direct H3K4 methylation. Nat Commun 13, 4521 (2022).

65. S. Chen, Y. Zhou, Y. Chen, J. Gu, fastp: an ultra-fast all-in-one FASTQ preprocessor. Bioinformatics 34, i884–i890 (2018).

66. B. Langmead, S. L. Salzberg, Fast gapped-read alignment with Bowtie 2. Nat Methods 9, 357–359 (2012).

67. P. Toolkit, Broad Institute.[Internet]. Available in: http://broadinstitute.github.io/picard, (2019).

68. P. Danecek et al., Twelve years of SAMtools and BCFtools. Gigascience 10, (2021).

69. Y. Zhang et al., Model-based analysis of ChIP-Seq (MACS). Genome Biol 9, R137 (2008).

70. S. Heinz et al., Simple combinations of lineage-determining transcription factors prime cis-regulatory elements required for macrophage and B cell identities. Mol Cell 38, 576–589 (2010).

71. F. Ramírez et al., deepTools2: a next generation web server for deep-sequencing data analysis. Nucleic Acids Res 44, W160–165 (2016).

72. T. L. Bailey, J. Johnson, C. E. Grant, W. S. Noble, The MEME Suite. Nucleic Acids Res 43, W39–49 (2015).

73. T. Nobusawa et al., Extranuclear function of Arabidopsis HOOKLESS1 regulates pleiotropic developmental processes in a non-cell-autonomous manner. New Phytol 246, 616–630 (2025).

74. J. Schindelin et al., Fiji: an open-source platform for biological-image analysis. Nat Methods 9, 676-682 (2012).

75. T. Romeis, A. A. Ludwig, R. Martin, J. D. Jones, Calcium-dependent protein kinases play an essential role in a plant defence response. EMBO J 20, 5556–5567 (2001).

