## Supplementary file for "Osmotic-stress-inducible nuclear condensates restrict gene inducibility"

**Supplementary Materials**

### Materials and Methods

#### Plant materials and growth conditions

*Arabidopsis thaliana* (Arabidopsis) Heynh. ecotype Columbia plants were grown on germination medium (GM) agar plates (half-strength Murashige and Skoog media (*48*) containing 0.83% (w/v) Bacto Agar, 3% (w/v) sucrose, 0.05% (w/v) MES, 1×Gamborg's B5 vitamin solution, pH5.7 (KOH)) at 22°C under a 16-h photoperiod at a photon flux density of 100 μmol $m^{-2}$ $sec^{-1}$ as described previously (*49*). Sterilized seeds with 70% ethanol and bleach were incubated at 4°C under dark conditions for 2 days for stratification. The 16-d-old plants were transferred to vermiculite containing soil (Asahi Industry, Japan) and grown under similar conditions. The T-DNA insertion lines of *NCED3* (*nc3-2*) and the transgenic plants expressing a nuclear marker (pro*RPS5a*:*H2B-mRuby*) was described previously (*27, 50*). Transgenic plants were generated by the floral dip method (*51*) using *Agrobacterium tumefaciens* (Agrobacterium) GV3101 (pMP90) cells. The CRISPR-Cas9 mutants were generated as previously described (*52*). Briefly, the pDe-CAS9 vector which harbors sgRNAs targeting *ALBA4/5/6* was transformed and selected the T1 plants by hygromycin and claforan. Genomic mutations were detected using the T2 plants by Sanger sequencing and, the CRISPR-Cas9-free plants were isolated in the T3 generation. The primers used to detect the T-DNA and CRISPR-Cas9-mediated mutations are shown in table S13.

#### Plasmid construction

To generate the constructs for iChIP (*21*), the *GAL4-BD* and *GFP-nls* coding sequences and the *35S* promoter region were inserted into pGreen0129 vector (*53*) between the *Pst*I-*Sal*I, *Sal*I-*Apa*I, and the *Not*I-*Spe*I sites, respectively. The 2.3-kb *NCED3* promoter region was inserted into pGK-GUS vector (*13*) between *Spe*I-*Sal*I sites. The GAL4 upstream activating sequence

(UAS) was inserted into *Xba*I-*Spe*I sites, and repeated UAS was made by excision of UAS at *Not*I-*Spe*I sites and insertion of the excised UAS into *Not*I-*Xba*I sites of the *UAS*-pro*NCED3*:*GUS* pGK vector.

For generating CRISPR-Cas9 knockout mutants, to express multiple sgRNAs, the *Nhe*I and *Hind*III sites were introduced on the 5' and 3' end of the expression cassette of the pEn-Chimera vector (*52*) by PCR and recombination (pEn-Chimera-NH vector). Next, annealed primers to express sgRNAs targeting *ALBA4*, *ALBA5* and *ALBA6* were introduced into the *Bbs*I sites. The expressing cassette of sgRNA was excised by *Nhe*I-*Hind*III, and tandemly fused into the *Xba*I-*Hind*III sites. The multiple expressing cassettes of sgRNAs were introduced into the pDe-Cas9 vector by recombination.

For GUS staining, the coding sequence of *GUSplus* was inserted into the *Spe*I-*Pst*I sites of the pB7WG2 vector without the cassette of the LR reaction (*54*) (GUS pB7WG2 vector) by NEBuilder® HiFi DNA Assembly Cloning Kit (New England Biolabs, Japan). Afterward, the promoter regions of *ALBA4*, *ALBA5* and *ALBA6* were inserted into *Kpn*I-*Apa*I sites by NEBuilder® HiFi DNA Assembly Cloning Kit.

For complementation assays, the coding sequence of *GFP* and *GFP-nls* was inserted into the *Spe*I-*Pst*I sites of the pB7WG2 vector without the cassette of the LR reaction (*54*) (GFP pB7WG2 and GFP-nls pB7WG2 vectors) by NEBuilder® HiFi DNA Assembly Cloning Kit. Afterward, the genomic sequences of *ALBA4* and *ALBA5* including the promoter regions (pro*ALBA4*:g*ALBA4* or pro*ALBA5*:g*ALBA5*) were inserted into the *Kpn*I-*Eco*RV sites of the GFP pB7WG2 or GFP-nls pB7WG2 vectors by NEBuilder® HiFi DNA Assembly Cloning Kit. To complement the deleted ALBA4, the promoter region of *ALBA4* was inserted into the *Kpn*I-*Apa*I sites of the GFP pB7WG2 vector. Afterward, the coding sequence of *ALBA4ΔIDR* was inserted into the *Apa*I-*Eco*RV sites by NEBuilder® HiFi DNA Assembly Cloning Kit.

To observe subcellular localization in *Nicotiana benthamiana*, the constitutive *35S* promoter region was inserted into the *Kpn*I-*Bam*HI sites of the GFP pB7WG2 vector above by

NEBuilder® HiFi DNA Assembly Cloning Kit. Afterward, the coding sequences of full length *ALBA4* into the *Xba*I-*Eco*RV sites. To observe the deleted *ALBA4*, *35S* promoter region was inserted into the *Kpn*I-*Bam*HI sites of the pB7WG2 vector without the cassette of the LR reaction (*54*) (pro35S pB7WG2 vector) by NEBuilder® HiFi DNA Assembly Cloning Kit. Afterward, the coding sequences of *ALBA4ΔIDR-GFP* and *ALBA4ΔN-GFP* were inserted into the *Bam*HI-*Pst*I sites. For a nuclear marker, the coding sequence of *tdTomato-nls* was inserted into the *Spe*I-*Pst*I sites of the pro35S pB7WG2 vector by NEBuilder® HiFi DNA Assembly Cloning Kit. For the PB and SB markers, the coding sequences of *RFP* was inserted into the *Spe*I-*Pst*I sites of the pro35S pB7WG2 vector. Afterward, the coding sequences of *DCP1* and *RBP47B* were inserted into the *Bam*HI-*Spe*I sites by NEBuilder® HiFi DNA Assembly Cloning Kit.

For *in vitro* protein synthesis, the coding sequence of *ALBA4-GFP* was inserted into the pCold TF vector (Takara, Japan) with deletion of the *TF* coding sequence by PCR with NEBuilder® HiFi DNA Assembly Cloning Kit.

The primers used for plasmid construction are shown in table S13.

**Stress treatment**

Dehydration treatments were exerted to plants as previously described (*13*), and 10-d-old plants grown on solid GM agar plates were transferred on parafilm. For the salt stress treatments, 10-d-old plants on grown on solid GM agar plates were transferred into liquid GM medium overnight for acclimation and were transferred into liquid GM medium supplemented with NaCl. Salt stress tolerance tests were performed as previous described (*55*), and 10-d-old plants grown on solid GM agar plates were transferred on GM agar plates supplemented with or without NaCl. Germination assays were performed as previously described (*54*). Sterilized seeds were incubated at 4°C under dark conditions for 3 days for stratification, and root protrusion was observed under a microscope (MICRONET, Japan) at different times during germination. Experiments were done in at least three biological replicates. Treatment of cordycepin was performed as described previously (*56*) with minor modifications. 10-d-old plants grown on solid GM agar plates were liquid GM medium overnight for acclimation and were transferred into liquid GM medium supplemented with NaCl. The plants were incubated for 12 hours, and transferred into liquid GM medium supplemented with 0.6 mM cordycepin (Fujifilm, Japan) and with and without NaCl. After vacuum infiltration for 1 min, the plants were incubated in the liquid media with and without NaCl.

**RIKEN Integrated Plant Phenotyping System (RIPPS)**

Plant phenotypes under drought stress on soil were analyzed using the RIPPS, an automated phenotyping platform (*25*), which can regulate soil moisture to any level with automated weigh and watering system. Six-day old seedlings grown on soil-filled plates were transferred to pots filled with standard potting mixture (Professional-baido, Daio Chemical, Tokyo, Japan). The soil water content (SWC) of the pots was adjusted every 2 h to 0.24 SWC (g $H_2O$ $g^{-1}$ dry soil) for the control treatments. For drought treatments, SWC was adjusted to 0.24 SWC for 7 days and then set to 0.1 SWC. To evaluate plant growth, projected leaf area was calculated from color images obtained with a color camera (Grasshopper3 9.1 MP Color USB3 Vision, Teledyne FLIR LLC, Hudson, NH, USA) using originally developed imaging software (LPIXEL Inc., Tokyo Japan). Thermal images were obtained using an infrared camera (FLIR T640, Teledyne FLIR LLC, Hudson, NH, USA). Leaf temperature during the daytime was calculated using originally developed imaging software that calculates the average temperature of plants by overlaying a mask image of plant parts extracted from a color image onto a thermal image (LPIXEL Inc., Tokyo Japan).

**Insertional chromatin immunoprecipitation (iChIP)**

The screen of iChIP was performed as previously described (*21*) with modifications. Two grams of seedlings after dehydration stress for 3 hours were submerged in 25 ml of 1% paraformaldehyde and vacuum infiltrated for 2 min, and incubated at room temperature for 30 min. The crosslinking was stopped by adding 3 ml of 1.25 M glycine. The seedlings were vacuum infiltrated for 2 min, and incubated at room temperature for 10 min. The crosslinked seedlings were rinsed with distilled water, and ground to a fine powder with mortars and pestles. The samples were resuspended in 25 ml of Extraction buffer 1 (0.4 M sucrose, 10 mM Tris-HCl pH 8.0, 5 mM β-Mercaptoethanol, 0.1 mM PMSF, 1×Complete EDTA-free), and filtered with two layers of Miracloth. The filtered samples were centrifuged at 2,880 *g* for 15 min at 4°C. The supernatant was removed, and the pellet was resuspended in 1 ml of Extraction buffer 2 (0.25 M sucrose, 10 mM Tris-HCl pH 8.0, 10 mM $MgCl_2$, 1% TritonX-100, 5 mM β-Mercaptoethanol, 0.1 mM PMSF, 1×Complete EDTA-free). The resuspended samples were transferred to 1.5 ml tubes and incubated on ice for 5 min. The samples were centrifuged at 14,000 *g* for 10 min at 4°C, and the supernatant was removed. The pellet was resuspended in 300 µl of Extraction buffer 3 (1.7 M sucrose, 10 mM Tris-HCl pH 8.0, 2 mM $MgCl_2$, 0.15% TritonX-100, 5 mM β-Mercaptoethanol, 0.1 mM PMSF, 1×Complete EDTA-free). The solutions were centrifuged at 14,000 *g* for 60 min at 4°C. The supernatant was removed, and the pellet was resuspended in 1 ml of Nuclei lysis buffer (10 mM Tris-HCl pH 8.0, 1 mM EDTA, 0.5 M NaCl, 1% Triton X-100, 0.5% sodium deoxycholate, 0.5% lauroylsarcosine, 1×Complete EDTA-free), and incubated on ice for 10 min. The solutions were centrifuged at 4,000 *g* for 10 min at 4°C. The supernatant was removed, and the pellet was resuspended in 400 µl of MLB3 (10 mM Tris-HCl pH 8.0, 1 mM EDTA, 0.5 mM EGTA, 150 mM NaCl, 0.1% sodium deoxycholate, 0.1% SDS, 1×Complete EDTA-free). The chromatin was sonicated by Bioruptor UCD-250 (Cosmo Bio, Japan) (15 s -on, 30 sec -off cycle, 15 min 200 W). The sonicated chromatin was centrifuged at 13,000 *g* for 10 min at 4°C. The supernatant was transferred into a new tube, and added with 100 µl of MLB-T (MLB3 + 5% Triton X-100). The GFP-Trap Magnetic Particles M-270 (Proteintech, Japan) were added to the solution, and rotated at 4°C overnight. The supernatant was removed on a magnet stand, and the beads were washed with Low salt wash buffer (20 mM Tris-HCl pH8.0, 150 mM NaCl, 0.1% SDS, 1% Triton X-100, 2 mM EDTA) twice, High salt wash buffer (20 mM Tris-HCl pH8.0, 500 mM NaCl, 0.1% SDS, 1% Triton X-100, 2 mM EDTA) twice, LiCl wash buffer (10 mM Tris-HCl pH8.0, 0.25 M LiCl, 1% NP40, 1% sodium deoxycholate, 1 mM EDTA) twice, and TE buffer once. The proteins were eluted with 40 µl of SDS sample buffer with incubation at 98°C for 30 min. Mass spectrometry was performed as described previously (*57*).

**RNA extraction and quantitative PCR (qPCR)**

The total RNA from seedlings was extracted using Monarch® Total RNA Miniprep Kit (New England Biolabs, Japan) according to the supplier's instruction. The complementary DNA (cDNA) was synthesized using Verso cDNA Synthesis Kit (Thermo Fisher Scientific, Japan) according to the supplier's instruction using the Random Hexamer, and qPCR was performed using Takara Thermal Cycler Dice® Real Time System III (Takara, Japan) and Luna® Universal qPCR Master Mix (New England Biolabs, Japan) according to the supplier's instructions. Triplicate measurements were made for each cDNA sample, and the values were normalized according to the amounts of *ACT2*. Reproducibility was confirmed in multiple biological replicates. The primers used for qPCR are shown in table S13.

**RNA sequencing (RNA-seq) and data analysis**

Total mRNA was isolated from total RNA using the NEBNext Poly(A) mRNA Magnetic Isolation Kit (New England Biolabs, Japan), and cDNA libraries were constructed using the NEBNext Ultra II RNA Library Prep Kit for Illumina (New England Biolabs, Japan). Libraries were sequenced on NovaSeq 6000 platform (Illumina) as paired-end through Takara RNA sequencing services. Reads were first quality checked with FastQC v0.11.9 and then trimmed using Trimmomatic (*58*) v0.39 using options MINLEN:60 HEADCROP:5 CROP:50. STAR

(*59*) v2.7.10a with basic options was used to map trimmed reads to the Araport11 genome annotation downloaded from https://plants.ensembl.org/. Mapped reads were assigned to genomic features using featureCounts (*60*) v2.0.0 with default settings. The assembled raw count matrices were imported into R and analyzed with the DESeq2 package (*61*) v1.34.0. Genes with a sum of reads < 10 across all samples were removed prior to principal component and differential expression analysis. Gene ontology (GO) analysis was performed using Shiny GO ver 0.73.3 (*62*). *Cis* element enrichment analysis was performed as previously described (*63*) using 1-kb upstream sequences from the translational start sites obtained through Phytozome Version 14 (https://phytozome-next.jgi.doe.gov/).

**Chromatin immunoprecipitation (ChIP)-qPCR**

ChIP assays were performed as previously described (*64*) with minor modifications. One gram of plant seedlings was frozen in liquid nitrogen, and ground to a fine powder with motors and pestles. The powdery samples were transferred to 25 ml of Nuclear isolation buffer (8.8 mM HEPES pH 7.6, 0.6 M sucrose, 4.4 mM KCl, 4.4 mM MgCl2, 4.4 mM EDTA pH 8.0, 0.3% (w/v) igepal, 0.1% (v/v) β-mercaptoethanol, 0.5 mM spermidine, 0.2 mM spermine, 1.5 mM EGS, 1% (v/v) formaldehyde, 1 mM pefabloc, 1×Complete EDTA-free). The resuspended sample was cross-linked with rotation for 10 min at room temperature, and cross-linking was stopped by adding 1.5 ml of 2 M glycine with rotation for 5 min at room temperature. The cross-linked sample was filtered with a 100 μm and following a 40 μm mesh cell strainer (AS ONE, Japan). The filtered solution was centrifuged at 3,000 *g* for 10 min at 4°C, and the supernatant was removed. The pellet was resuspended with 300 μl of Nuclear resuspend buffer (10 mM HEPES pH 7.6, 1 M sucrose, 5 mM KCl, 5 mM MgCl2, 5 mM EDTA pH 8.0), and carefully layered on 500 μl of Nuclear separation buffer (10 mM HEPES pH7.6, 1 M sucrose, 5 mM KCl, 5 mM MgCl2, 5 mM EDTA pH 8.0, 15% Percoll) in a 1.5 ml tube. The layered sample was centrifuged at 3,000 *g* for 5 min at 4°C, and the supernatant was removed. The pellet was resuspended with 1 ml of 1×RIPA buffer (50 mM Tris-HCl pH7.8, 0.15 M NaCl, 1 mM EDTA pH 8.0, 0.1 % SDS, 0.1 % Sodium deoxycholate, 1×Complete EDTA-free), and sonicated with the Covaris® M220 Focused-ultrasonicator™ (Covaris) using a Covaris milliTUBE 1 ml AFA Fiber (Treatment 1: Peak Incident Power: 50 W, Duty Factor: 10%, Cycles per burst: 200, Treatment time: 60 s and Treatment 2: Peak Incident Power: 75 W, Duty Factor: 10%, Cycles per burst: 200, Treatment time: 60 s, 17 cycles of Treatment 1 and 2, at 6°C). The sonicated sample was transferred to a 1.5 ml tube, and 50 μl of 20% Triton X-100 was added. The sample was centrifuged at 13,000 *g* for 3 min at 4°C, and 980 μl of the supernatant was transferred to a 1.5 ml tube. Additional 9.8 μl of 100×Complete EDTA-free stock solution was input to the sample, and 50 μl of the sample was used as "input". Immunoprecipitation (IP) was performed by adding 1 μl of the anti-GFP polyclonal antibody (Abcam, Japan) to the remaining sample with rotation at 4°C overnight. The Dynabeads™ Protein G (Thermo Fisher Scientific, Japan) (60 μl for each sample) were washed with 1 ml of PBS twice and 1 ml of 1×RIPA buffer once. The washed Dynabeads™ were resuspended in 60 μl of 1×RIPA buffer for each sample. The resuspended Dynabeads™ was added to the sample with the anti-GFP antibody, and incubated for 3 hours at 4°C with rotation. After the incubation, the tube was placed on a magnetic rack for 1 min and the supernatant was removed. The beads were washed by 1 ml of 1×RIPA buffer, 1 ml of medium RIPA buffer (50 mM Tris-HCl pH7.8, 0.3 M NaCl, 1 mM EDTA pH 8.0, 0.1 % SDS, 0.1 % Sodium deoxycholate, 1×Complete EDTA-free), 1ml of 2×RIPA buffer (100 mM Tris-HCl pH7.8, 0.3 M NaCl, 2 mM EDTA pH 8.0, 0.2 % SDS, 0.2 % Sodium deoxycholate, 1×Complete EDTA-free) and TE buffer with rotation for 10 min at 4°C. For elution, 100 μl of ChIP elution buffer (50 mM Tris-HCl pH7.5, 10 mM EDTA, 1 % SDS) was added, and the sample with beads was incubated for 15 min at 65°C. Reverse cross-linking was performed by incubating the sample at 65°C for 9 hours with 4 μl of 20 mg/ml Proteinase K enzyme (New England Biolabs, Japan). The 50 μl of input samples was also incubated similarly with 1 μl of the Proteinase K enzyme. After reverse cross-linking, the tube was placed on the magnetic rack for 1 min, and the supernatant was transferred to a 1.5 ml tube. The immunoprecipitated DNA was purified using the Monarch® PCR & DNA Cleanup Kit (5 μg) (New England Biolabs, Japan) according to the supplier's instruction. Concentrations of the purified DNA were measured using the Qubit dsDNA High Sensitivity Assay Kit (Thermo Fisher Scientific, Japan). ChIP-qPCR was performed using Takara Thermal Cycler Dice® Real Time System III (Takara, Japan) and Luna® Universal qPCR Master Mix (New England Biolabs, Japan) according to the supplier's instructions. Measurements of total six replicates using two biological replicates were made for each sample, and the values were normalized according to the amounts of input. The primers used for qPCR are shown in table S13.

**ChIP-sequencing (ChIP-seq) and data analysis**

ChIP-seq libraries were synthesized using Takara ThruPLEX® DNA-Seq Kit (Takara, Japan) according to the supplier's instruction. Sequencing of the ChIP-seq libraries was performed using the Novaseq 6000 (Illumina) with a 150 bp paired-end configuration. An *Arabidopsis thaliana* reference genome and transcript models of TAIR10 were downloaded from Ensemble. To remove the adapter sequences, the raw reads were trimmed using fastp version 0.23.4 (*65*). The trimmed ChIP-Seq reads were aligned to the reference genome TAIR10 using bowtie2 version 2.3.5.1 (*66*). Multi mapped and duplicate reads were removed using picard version 2.26.11 (*67*) and samtools version 1.15 (*68*). The retained reads were used to analyze the peak calling by macs2 version 2.2.7.1 (*69*). The peaks were annotated and enriched cis elements within the peaks were identified using homer version 4.7.2 (*70*). Correlation plots and Metagene plots were made by plotCorrelation and plotHeatmap, respectively, from deeptools version 3.5.5 (*71*). The enriched cis elements within 1 kb-promoter and 1 kb-downstream regions were identified through MEME Suite ver 5.5.9 (*72*) using the all 1 kb-promoter and 1 kb-downstream regions of Arabidopsis genes as controls.

**Quantification of plant hormones**

Plant hormones were quantified using LC-MS/MS as described previously (*73*), with modifications. Briefly, 100 mg of fresh tissue was suspended in 1 ml of 80% (v/v) methanol containing internal standards (2 ng each of $D_6$-ABA, $D_5$-IAA and $^{13}C_6$-JA-Ile; 0.5 ng of $D_6$-iP; Olchemim Ltd, Olomouc, Czech Republic). After homogenization, the extract was purified using a Bond Elut C18 column (Agilent Technologies, Santa Clara, CA, USA) and concentrated. The residue was resuspended in 100 μl of 50% methanol, and 1–2 μl was injected into an Agilent 1200 HPLC system with an Agilent 6460 triple quadrupole mass spectrometer. Separation was performed on a ZORBAX Eclipse XDB-C18 column (50×2.1 mm, 1.8 μm; Agilent Technologies). Plant hormones were monitored using multiple reaction monitoring (MRM), and were analyzed with MASSHUNTER software (Agilent Technologies). Plant hormone levels were calculated based on the ratio of analyte peak areas to stable isotope-labeled internal standards.

**GUS staining**

GUS staining was performed as described previously (*55*) with modifications. Plant seedlings were incubated 90% (w/w) acetone for 15 min on ice, and washed with 100 mM phosphate buffer. The seedlings were soaked in a Staining solution containing 0.26 mg/mL 5-bromo-4-chloro-3-indolyl β-D-glucuronide, 50 mM phosphate buffer, pH 7.0, and 20%(v/v) methanol at 28°C for 24 h. Pictures were taken by a stereomicroscope (SMZ18, Nikon, Tokyo, Japan) and NIS-Elements BR software (Nikon, Tokyo, Japan).

**Confocal image and imaging analysis**

Fluorescent protein signal was observed using an Olympus FV1200 confocal microscope with a UPLSAPO20X (NA = 0.75, WD = 0.6 mm) or UPLSAPO60XW (NA = 1.2, WD = 0.28) objective lens (Olympus, Tokyo, Japan) and FV10-ASW software ver. 4.2a (Olympus, Tokyo, Japan). Excitation lasers 473 nm and 559 nm and band pass filters 490-540 nm and 575-675 nm were used to detect the GFP and RFP/mRuby/tdTomato signals, respectively. The images were processed, and evaluation of condensates formation was performed by Fiji software (*74*). Fluorescence recovery after photobleaching (FRAP) experiments were performed through the FRAP mode of the FV10-ASW software ver. 4.2a to bleach a condensate in nuclei *in planta* or *in vitro* (488 nm, 100% intensity), and the recovery phase was imaged in time-series. For line scan analysis, a straight line in the picture was drown across condensates as the region of interest (ROI), and the intensity of the ROI was measured using Fiji.

**Synthesis and purification of recombinant proteins**

The His-ALBA4-GFP protein in *Escherichia coli* (*E. coli*) Rosetta(DE3) cells (Merck, Japan) was expressed in 100 ml of Overnight Express™ Instant LB Medium (Merck, Japan) according to the supplier's instruction. The protein was purified using TALON® Metal Affinity Resin (Clontech, Japan) and TALON® 2 ml Disposable Gravity Column (Clontech, Japan) according to the supplier's instruction. The protein concentration was measured by a Bradford protein assay (APRO, Japan) according to the supplier's instruction. For observation of ALBA4-GFP in *E. coli*, the *E. coli* cells after protein induction was incubated in the Overnight Express™ Instant LB Medium with and without NaCl. For observation of ALBA4-GFP condensates *in vitro*, Polyethylene Glycol 6000 (Fijifillm, Japan) was added to the purified ALBA4-GFP.

**Transient expression in *Nicotiana benthamiana* (tobacco) leaves**

Transient expression of proteins in tobacco leaves were performed as previously described (*75*). Briefly, Agrobacterium GV3101 (pMP90) cells harboring plasmids for protein expression were cultured at 28°C and then harvested by centrifugation. The cells were resuspended in infiltration buffer (10 mM $MgCl_2$, 10 mM MES, pH 5.6, and 150 mM acetosyringone) with an adjusted OD600 to 0.5. After incubation for 2 h at room temperature, the resuspended cells were infiltrated into leaves using a syringe. Five days after infiltration, expressed fluorescent proteins were observed under a confocal microscopy.

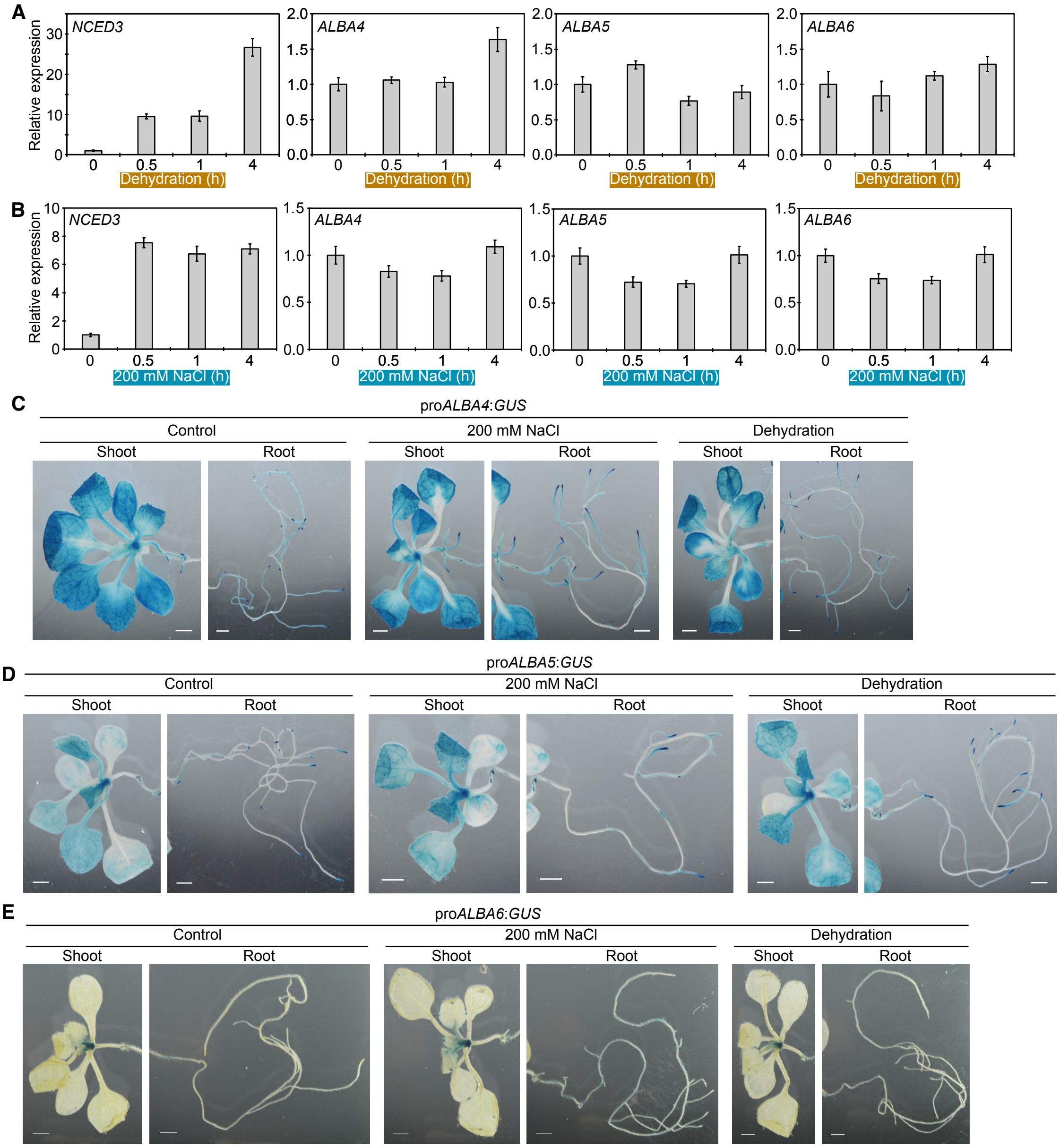


**Fig. S1. Expression patterns of *ALBA4*, *ALBA5* and *ALBA6* under osmotic stress conditions.** (**A** and **B**) Expression levels of *NCED3*, *ALBA4*, *ALBA5* and *ALBA6* under dehydration (A) and salt stress (B) conditions. Error bars indicate SD from triplicate technical repeats. (**C** to **E**) GUS staining of shoot and root tissues in the pro*ALBA4:GUS* (C), pro*ALBA5:GUS* (D) and pro*ALBA6:GUS* (E) plants at 14 d after sowing under control, dehydration and salt stress conditions (Scale bars: 1 mm).

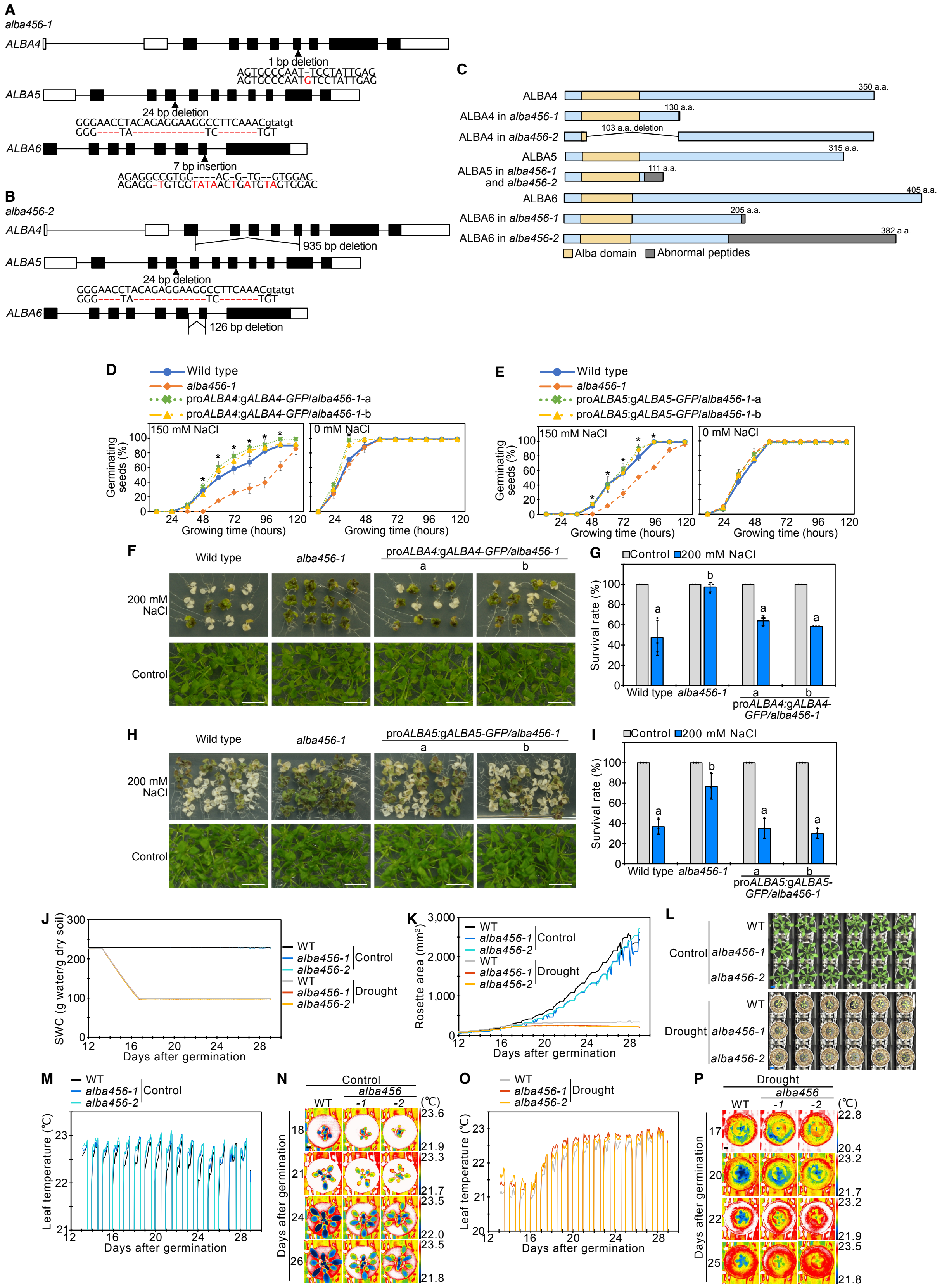
A
alba456-1
ALBA4
1 bp deletion
AGTGCCCAAT-TCCTATTGAG
AGTGCCCAATGTCCTATTGAG
ALBA5
24 bp deletion
GGGAACCTACAGAGGAAGGCCTTCAAACgtatgt
GGG----TA-------------TC-------TGT
ALBA6
7 bp insertion
AGAGGCCGTGG----AC-G-TG--GTGGAC
AGAGG-TGTGGTATAACTGATGTAGTGGAC
B
alba456-2
ALBA4
935 bp deletion
ALBA5
24 bp deletion
GGGAACCTACAGAGGAAGGCCTTCAAACgtatgt
GGG----TA-------------TC-------TGT
ALBA6
126 bp deletion
C
ALBA4
350 a.a.
ALBA4 in alba456-1
130 a.a.
ALBA4 in alba456-2
103 a.a. deletion
ALBA5
315 a.a.
ALBA5 in alba456-1 and alba456-2
111 a.a.
ALBA6
405 a.a.
ALBA6 in alba456-1
205 a.a.
ALBA6 in alba456-2
382 a.a.
Alba domain
Abnormal peptides
D
Wild type
alba456-1
proALBA4:gALBA4-GFP/alba456-1-a
proALBA4:gALBA4-GFP/alba456-1-b
150 mM NaCl
0 mM NaCl
Germinating seeds (%)
Growing time (hours)
E
Wild type
alba456-1
proALBA5:gALBA5-GFP/alba456-1-a
proALBA5:gALBA5-GFP/alba456-1-b
150 mM NaCl
0 mM NaCl
Germinating seeds (%)
Growing time (hours)
F
Wild type
alba456-1
proALBA4:gALBA4-GFP/alba456-1
a
b
200 mM NaCl
Control
G
Control
200 mM NaCl
Survival rate (%)
Wild type
alba456-1
proALBA4:gALBA4-GFP/alba456-1
H
Wild type
alba456-1
proALBA5:gALBA5-GFP/alba456-1
a
b
200 mM NaCl
Control
I
Control
200 mM NaCl
Survival rate (%)
Wild type
alba456-1
proALBA5:gALBA5-GFP/alba456-1
J
SWC (g water/g dry soil)
Days after germination
WT
alba456-1
alba456-2
Control
Drought
K
Rosette area (mm²)
Days after germination
L
Control
Drought
WT
alba456-1
alba456-2
M
Leaf temperature (℃)
Days after germination
N
Control
alba456
WT
-1
-2
(℃)
Days after germination
O
Leaf temperature (℃)
Days after germination
P
Drought
alba456
WT
-1
-2
(℃)
Days after germination

**Fig. S2. Phenotypic analysis of the complemented lines of ALBA4 and ALBA5 into the *alba456-1* triple mutant, and phenotypic analysis of the *alba456* triple mutants using RIPPS.** (**A** and **B**) Gene models of *ALBA4*, *ALBA5* and *ALBA6* and the mutations introduced by CRISPR-Cas9 editing in the two lines of *alba4 alba5 alba6* (*alba456*) triple mutants, *alba456-1* (A) and *alba456-2* (B). The coding regions, untranslated regions, and introns are indicated as a closed box, open boxes, and lines, respectively. (**C**) Schematic models of translated ALBA4, ALBA5 and ALBA6 proteins in wild type, *alba456-1* and *alba456-2* mutants. The conserved Alba motifs and abnormal peptide sequences are shown in yellow and gray boxes. ) (**D** and **E**) Germination assays in wild type, *alba456-1* and the complemented lines with ALBA4 (pro*ALBA4*:g*ALBA4-GFP*/*alba456-1*-a, b) (D) or ALBA5 (pro*ALBA5*:g*ALBA5-GFP*/*alba456-1*-a, b) (E). Error bars indicate SD from three biological replicates. Asterisks indicates significant differences between *alba456-1* and the complemented lines ($P < 0.05$, Tukey's multiple range test). (**F** to **I**) Salt stress tolerance tests in wild type, *alba456-1* and the complemented lines with ALBA4 (pro*ALBA4*:g*ALBA4-GFP*/*alba456-1*-a, b) (F and G) or ALBA5 (pro*ALBA5*:g*ALBA5-GFP*/*alba456-1*-a, b) (H and I). Images with and without salt stress (F and H) and the survival ratios (G and I) are shown. Error bars indicate SD from three biological replicates. Letters indicates significant differences from wild type ($P < 0.05$, Tukey's multiple range test). (**J**) Soil water content (SWC) under well-watered and drought stress conditions in RIPPS. (**K** and **L**) Rosette areas of wild type and *alba456* triple mutants under control and drought stress conditions in RIPPS. The transition of quantitative data of the rosette areas in wild type, *alba456-1* and *alba456-2* under control and drought stress conditions (K) and the images on the 28$^{th}$ day (L) are shown (Scale bars: 2 cm). (**M** to **P**) Leaf temperature of wild type and *alba456* triple mutants under control (M and N) and drought stress (O and P) conditions in RIPPS. Quantitative data (M and O) and images on several timepoint (N and P) are shown (Scale bars: 1 cm).

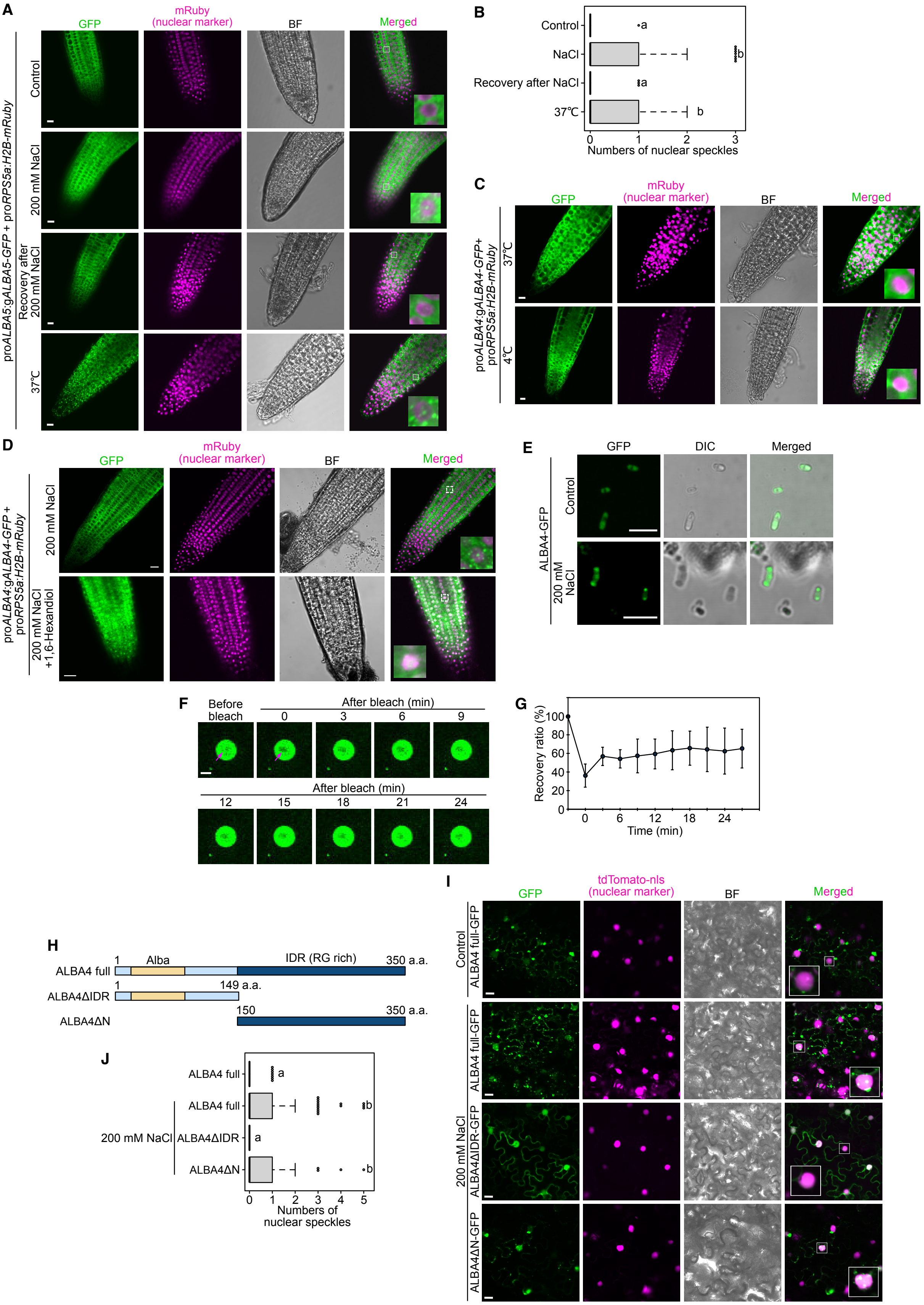
A
GFP
mRuby (nuclear marker)
BF
Merged
proALBA5:gALBA5-GFP + proRPS5a:H2B-mRuby
Control
200 mM NaCl
Recovery after 200 mM NaCl
37℃
B
Control
NaCl
Recovery after NaCl
37℃
a
b
a
b
0 1 2 3
Numbers of nuclear speckles
C
GFP
mRuby (nuclear marker)
BF
Merged
proALBA4:gALBA4-GFP+ proRPS5a:H2B-mRuby
37℃
4℃
D
GFP
mRuby (nuclear marker)
BF
Merged
proALBA4:gALBA4-GFP + proRPS5a:H2B-mRuby
200 mM NaCl
200 mM NaCl +1,6-Hexandiol
E
GFP
DIC
Merged
ALBA4-GFP
Control
200 mM NaCl
F
Before bleach
After bleach (min)
0 3 6 9
After bleach (min)
12 15 18 21 24
G
Recovery ratio (%)
100 80 60 40 20 0
0 6 12 18 24
Time (min)
H
ALBA4 full
1
Alba
IDR (RG rich)
350 a.a.
ALBA4ΔIDR
1
149 a.a.
ALBA4ΔN
150
350 a.a.
I
GFP
tdTomato-nls (nuclear marker)
BF
Merged
Control
ALBA4 full-GFP
200 mM NaCl
ALBA4 full-GFP
ALBA4ΔIDR-GFP
ALBA4ΔN-GFP
J
ALBA4 full
200 mM NaCl
ALBA4 full
ALBA4ΔIDR
ALBA4ΔN
a
b
a
b
0 1 2 3 4 5
Numbers of nuclear speckles

**Fig. S3. Analysis of molecular characteristics of ALBA condensates under abiotic stress conditions.** (**A** and **B**) Confocal microscopic analysis of ALBA5-GFP under stress conditions. Confocal images (A) and numbers of nuclear speckles under each condition (B) are shown. H2B-mRuby was expressed as a nuclear marker. Scale bars represent 10 μm. Letters indicates significant differences ($P$ < 0.05, Tukey's multiple range test). (**C**) Confocal microscopic analysis of ALBA4-GFP under heat and cold stress conditions. Scale bars represent 10 μm. (**D**) Treatment of 1,6-hexandiol on the ALBA4 condensates during salt stress (Scale bar: 10 μm). (**E**) Formation of ALBA4-GFP condensates under salt stress conditions in *Escherichia coli*. Scale bars represent 5 μm. (**F** and **G**) FRAP of ALBA4 nuclear condensates during salt stress *in vitro*. Time-lapse images (F) and plot showing the recovery after photobleaching (G) are shown. The red arrows indicate a bleached part (Scale bar: 2 μm). Error bars indicate SD from twelve independent experiments. (**H**) Schematic models of translated the ALBA4 protein. The conserved Alba motifs and IDR are shown. (**I** and **J**) Confocal microscopic analysis of full length and truncated regions of ALBA4-GFP transiently expressed in *Nicotiana benthamiana* leaves treated with and without salt stress. Confocal images (I) and numbers of nuclear speckles under each condition (J) are shown. tdTomato-nls was expressed as a nuclear marker. Scale bars represent 10 μm. Letters indicates significant differences ($P$ < 0.05, Tukey's multiple range test). Error bars indicate SD from triplicate technical repeats. Asterisks indicate significant differences at each time point ($P$ < 0.05, Tukey's multiple range test).

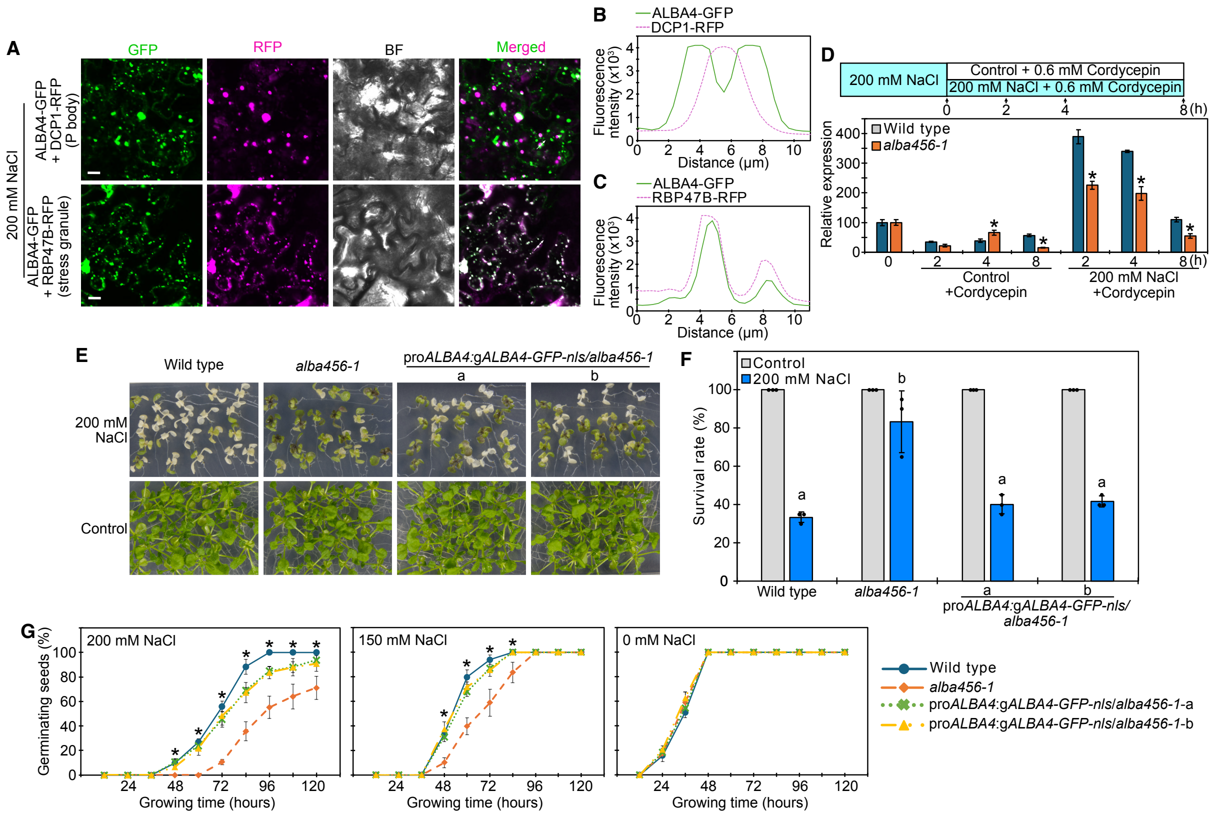


**Fig. S4. Analysis of ALBA4 condensates in cytosol and nuclei under salt stress conditions.** (**A** to **C**) Confocal microscopic analysis for colocalization of ALBA4 with a P body maker protein DCP1and a stress granule marker protein RBP47B under salt stress conditions. Confocal images (A) and line profiling of fluorescence intensities (B and C) are shown. Scale bars represent 10 μm. The white arrows in upper and bottom images represent planes for generating line profiles of ALBA4-GFP with DCP1-RFP (B) and RBP47B-RFP (C). (**D**) Decay profiles of the *NCED3* mRNA under control and salt stress conditions with treatment of Cordycepin. Error bars indicate SD from triplicate technical repeats. Asterisks indicates significant differences from wild type at each time point ($P < 0.05$, Student's t test). (**E** and **F**) Salt stress tolerance of the complemented lines of ALBA4-GFP-nls into the *alba456* triple mutant (pro*ALBA4*:g*ALBA4-GFP-nls*/*alba456-1*-a and b). Images with and without salt stress (E) and the survival ratios (F) are shown. Error bars indicate SD from three biological replicates. Letters indicates significant differences during salt stress ($P < 0.05$, Tukey's multiple range test). (**G**) Germination ratios of the pro*ALBA4*:g*ALBA4-GFP-nls*/*alba456-1* plants under slat stress conditions. Error bars indicate SD from six biological replicates. Asterisks indicates significant differences in both complemented lines from wild type at each time point ($P < 0.05$, Tukey's multiple range test).

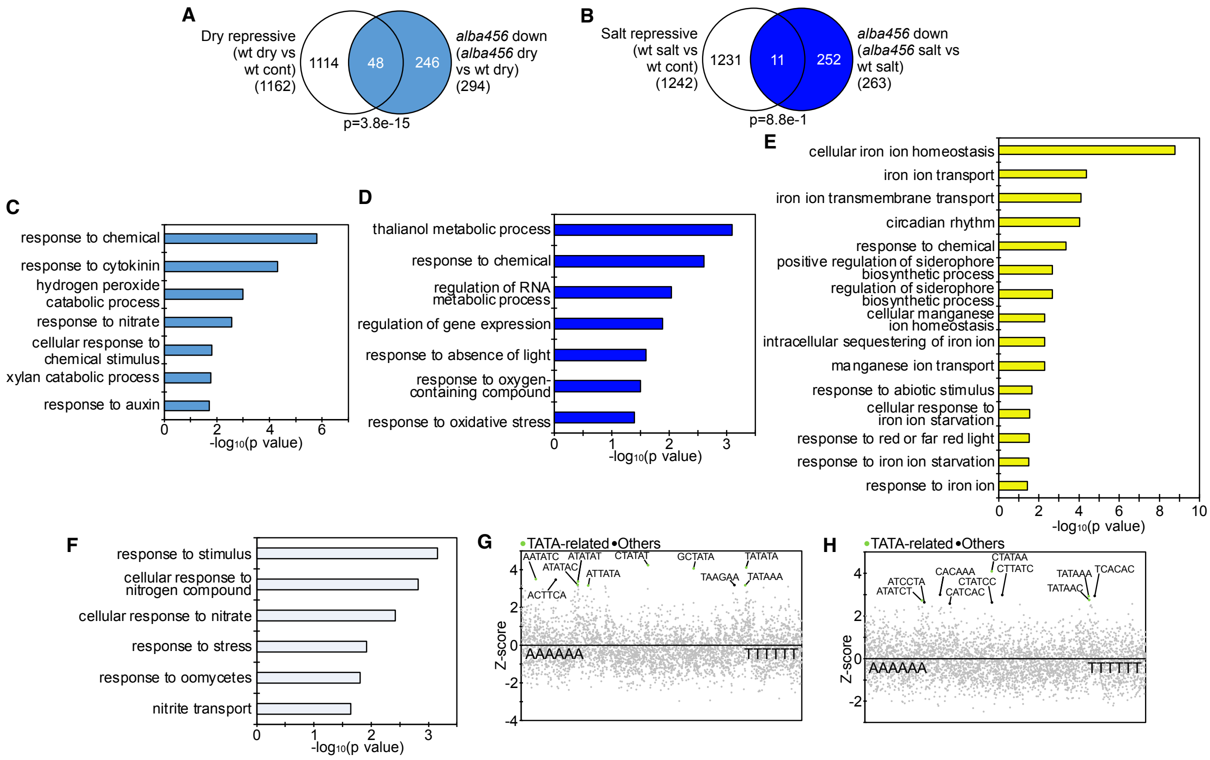


**Fig. S5. Transcriptomic analysis of *alba456-1* triple mutant under osmotic stress conditions.** (**A** and **B**) Venn diagrams showing overlaps between stress suppressive genes and downregulated genes in *alba456-1* during dehydration (A) and salt (B) stress. P values are calculated by Fisher's exact test. (**C** and **D**) Enrichment of GOs among downregulated genes in *alba456-1* during dehydration (C) and salt (D) stress. (**E** and **F**) Enrichment of GOs in *alba456-1* under control conditions using up- (E) and downregulated (F) genes ($|Log_2FC| \geq 1$, $p<0.05$). (**G** and **H**) Overrepresentation analysis of hexamer motifs in the 1-kb promoter regions of the top 200 downregulated genes in *alba456-1* during dehydration (G) and salt (H) stress. Z scores (y axis) for the observed frequencies of all hexamer motifs (x axis) are presented in the scatter plot. The TATA-related and other top 10 enriched motifs are highlighted, in green and black, respectively.

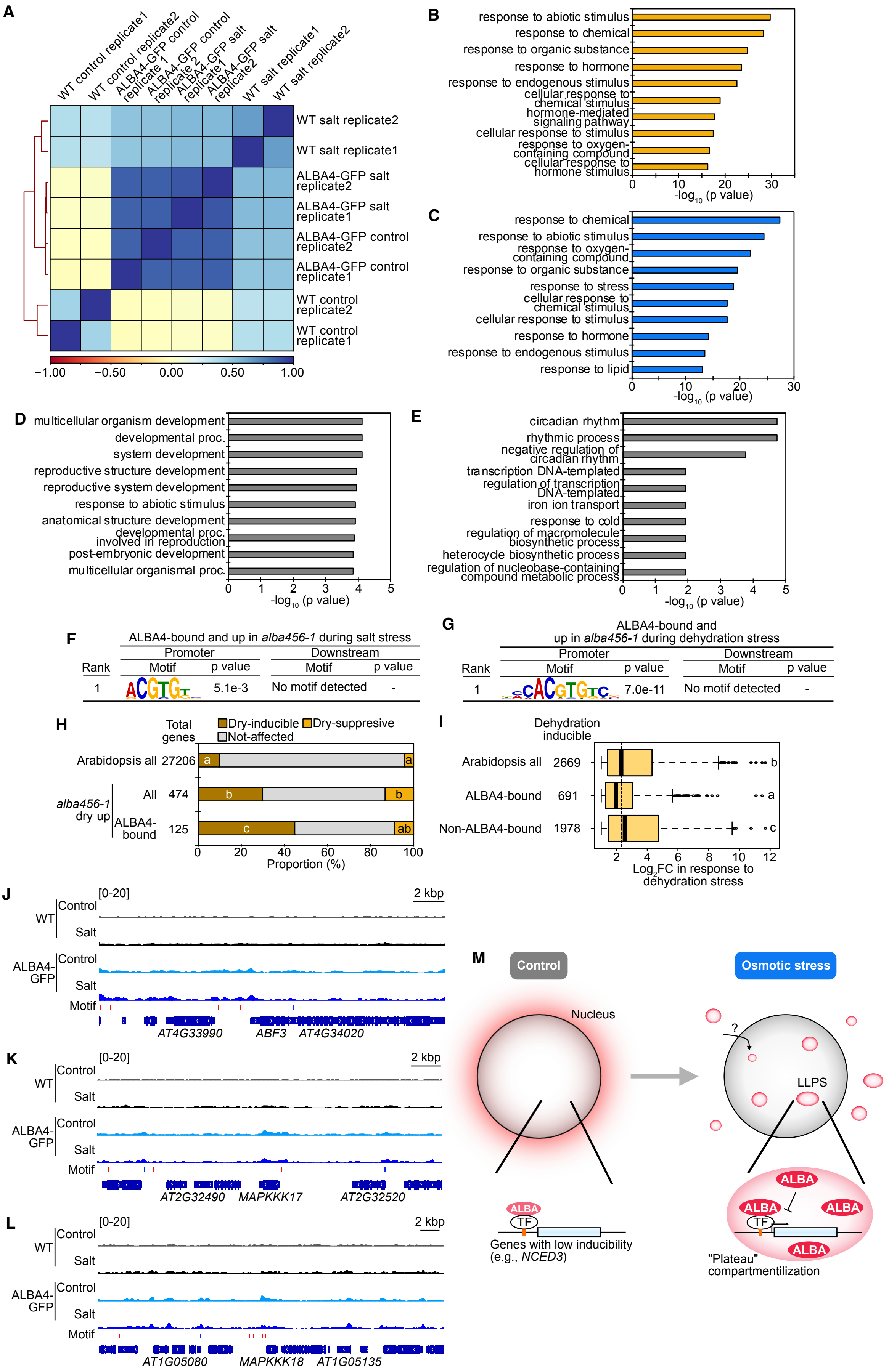
A
B
response to abiotic stimulus
response to chemical
response to organic substance
response to hormone
response to endogenous stimulus
cellular response to chemical stimulus
hormone-mediated signaling pathway
cellular response to stimulus
response to oxygen-containing compound
cellular response to hormone stimulus
C
response to stress
response to lipid
D
multicellular organism development
developmental proc.
system development
reproductive structure development
reproductive system development
anatomical structure development
developmental proc. involved in reproduction
post-embryonic development
multicellular organismal proc.
E
circadian rhythm
rhythmic process
negative regulation of circadian rhythm
transcription DNA-templated
regulation of transcription DNA-templated
iron ion transport
response to cold
regulation of macromolecule biosynthetic process
heterocycle biosynthetic process
regulation of nucleobase-containing compound metabolic process
F
ALBA4-bound and up in alba456-1 during salt stress
G
ALBA4-bound and up in alba456-1 during dehydration stress
H
I
J
K
L
M
Control
Osmotic stress
Nucleus
LLPS
ALBA
TF
Genes with low inducibility (e.g., NCED3)
"Plateau" compartmentilization

**Fig. S6. Integrative analysis of transcriptomic data in *alba456-1* and ChIP-seq of ALBA4.** (**A**) Correlation heatmap showing the strong correlation between two biological replicates and stress conditions of ALBA4 ChIP-seq data. (**B** to **D**) Enrichment of GOs among the ALBA4-bound genes under both control and salt (B), salt-specific (C) and control-specific (D) conditions. (**E**) Enrichment of GOs among the upregulated genes in *alba456-1* under control conditions which were bound by ALBA4 under control conditions. (**F** and **G**) Enriched motifs identified by MEME using the 1-kb upstream regions from translational start sites and 1-kb downstream regions from translational stop sites among the upregulated genes in *alba456-1* during salt (F) and dehydration (G) stress which were bound by ALBA4 comparing to the 1-kb upstream and downstream regions of Arabidopsis all genes, respectively. (**H**) Plots showing proportions of stress inducible genes among Arabidopsis all genes and upregulated genes in *alba456-1* during dehydration stress which were bound by ALBA4. Letters represent significant differences within the proportions of stress-inducible or suppressive genes ($P < 0.05$, pairwise Fisher's exact test with Benjamini–Hochberg correction). (**I**) Plots showing dehydration stress inducibility of genes bound by ALBA4. Letters represent significant differences ($P < 0.05$, pairwise Wilcoxon test with Benjamini–Hochberg correction). (**J** to **L**) IGV screenshots showing the ChIP-seq signals of ALBA4 on *ABF3* (J), *MAPKKK17* (K) and *MAPKKK18* (L). All tracks represent read density normalized to sequencing depth. The gene models are shown at the bottom with arrow heads showing the direction of transcription. Red and blue bars represent locations of G box (CACGTG) and GGGCCC motifs, respectively. (**M**) Schematic model of molecular mechanisms in which ALBAs suppress stress inducibility of osmotic stress-inducible genes. Under control conditions, ALBAs are mainly localized in cytosol meanwhile small amount of the ALBA proteins in nuclei are bound to the target genes including *NCED3*. Any transcription factors binding to G box may tether ALBAs to the promoter regions of the target genes. Upon osmotic stress conditions, formation of the ALBA condensates through LLPS is induced in both cytosol and nuclei. The ALBA condensates in nuclei function to suppress stress inducibility of the ALBA target genes such as *NCED3*. Nuclear translocation of the ALBA proteins from cytosol might trigger phase separation of ALBAs in nuclei.
